# Root morphology and exudate availability is shaped by particle size and chemistry in *Brachypodium distachyon*

**DOI:** 10.1101/651570

**Authors:** Joelle Sasse, Jacob S. Jordan, Markus DeRaad, Katherine Whiting, Katherina Zhalnina, Trent Northen

## Abstract

Root morphology and exudation define a plants sphere of influence in soils, and are in turn shaped by the physiochemical characteristics of soil. We explored how particle size and chemistry of growth substrates affect root morphology and exudation of the model grass *Brachypodium distachyon*. Root fresh weight and root lengths were correlated with particle size, whereas root number and shoot weight remained constant. Mass spectrometry imaging suggested that both, root length and number shape root exudation. Exudate metabolite profiles detected with liquid chromatography / mass spectrometry were comparable for plants growing in glass beads or sand with various particles sizes, but distinct for plants growing in clay. However, when exudates of clay-grown plants were collected by removing the plants from the substrate, their exudate profile was similar to sand- or glass beads-grown plants. Clay particles sorbed 20% of compounds exuded by clay-grown plants, and 70% of compounds of a defined exudate medium. The sorbed compounds belonged to a range of chemical classes, among them nucleosides/nucleotides, organic acids, sugars, and amino acids. Some of the sorbed compounds could be de-sorbed by a rhizobacterium (*Pseudomonas fluorescens* WCS415), supporting its growth. We show that root morphology is affected by substrate size, and that root exudation in contrast is not affected by substrate size or chemistry. The availability of exuded compounds, however, depends on the substrate present. These findings further support the critical importance of the physiochemical properties of soils are crucial to consider when investigating plant morphology, exudation, and plant-microbe interactions.

## Introduction

Plant roots shape their environment in various ways, and are in turn shaped by physiochemical properties of the surrounding soil. Roots shape soil by dislocating particles, by polymer production, and by release of a wide variety of compounds (root exudation). Root exudates alter pH and the chemical composition around roots. Overall, root presence in soils results in formation of larger soil aggregates and increases water-holding capacity (Six et al., 2004). The plant-induced changes in chemistry can lead to weathering of minerals (Uroz et al., 2015), and alter the composition of microbial communities (Carson et al., 2007).

Soils are often characterized by their particle size and mineralogy. Typical soil particles range from small (< 50 µm) to large (> 2 mm), determining physical parameters such as water binding capacity (Six et al., 2004; Six and Paustian, 2014; Rellán-Álvarez et al., 2016). Differently sized sandstones are associated with different microbial numbers, with small sandstones (2 mm) being more densely populated by microbes than larger rocks (Certini et al., 2004). Minerals differ in their structure (e.g. accessible surface), and in their surface charge, determining if and how they interact with dissolved compounds (Swenson et al., 2015a). Differences in mineralogy, nutrient content, and weathering of likely result in associations with distinct microbial communities (Carson et al., 2009; Uroz et al., 2015; Whitman et al., 2018).

An influence of particle size on plant morphology was shown in previous studies, where maize root and shoot growth was reduced in 1 mm glass beads vs. hydroponic conditions, the root:shoot ratio was elevated, and root morphology was altered (Veen, 1982; Boeuf-Tremblay et al., 1995; Groleau-Renaud et al., 1998). One study found a small reduction in sugar exudation and an increase in amino acid exudation (Boeuf-Tremblay et al., 1995), whereas another study found no change in carbon exudation (Groleau-Renaud et al., 1998), making the effect of particle size on exudation less clear than its effect on root morphology.

It is more evident that substrate chemistry affects exudation. Maize growing in stonewool for example exudes higher amounts of organic acids and sugars compared to maize growing in glass beads, possibly due to the presence of aluminum ions, which increase organic acids exudation (Kamilova et al., 2006). Activated charcoal was found to sorb most of the phenolic compounds exuded by Arabidopsis, altering the plants root morphology (Caffaro et al., 2011). Different soils resulted in distinct lettuce root weight and root lengths, as well as in distinct presence of some rhizosphere metabolites (Neumann et al., 2014). As this study was performed in soils with different chemistry, granule sizes, and microbial communities, it is not possible to identify the causal factor of the changes observed (Neumann et al., 2014; Schreiter et al., 2014). Overall, the effect of physical structure and chemistry of commonly found substrates such as sand and clay on root morphology and exudation are not well studied to date.

To determine how particle size and chemistry of the growth substrate shape root morphology and exudation, reductionistic experimental setups are desirable: A sterile environment permits a focus on plant metabolism without the additional layer of microbial metabolism present in a natural environment. The use of an inert substrate with defined particle size enables the investigation of physical effects on root morphology and exudation, whereas the use of a substrate with defined surface chemistry allows to study chemical interactions of exudates with particles.

Here, we chose to investigate the effect of particle size and chemistry on root morphology and exudation in *Brachypodium distachyon*. Specifically, we asked three questions: 1) whether the altered root morphology observed in maize growing in physically restricted conditions hold true in a model grass, 2) if and how the exudate profile changes with particle size, and 3) how root morphology and exudation are influenced by substrate chemistry. To answer these questions, *B. distachyon* was grown in glass beads with various sizes, as well as in sand and in clay particles. Root morphology was assessed in all conditions, as well as the exudate metabolic profile of plants growing in the substrate (‘*in situ’*), and subsequently of the plants transferred to a hydroponic solution (‘*in vitro’*), to be able to distinguish changes due to altered plant metabolism from changes due to interaction of exudates with substrates. Root weight and root length were found to correlate with particle size, whereas shoot weight and root number remained constant. Root exudation was correlated with root weight, and exudation profiles remained unchanged in the various conditions. Clay sorbed a large degree of exudates, altering the amount of exudates freely available around root, and clay-sorbed exudates supported growth of a rhizobacterium. Our results highlight the importance of taking into account soil structure and chemistry when studying plant – soil interactions.

## Results

### Metabolite sorption to substrates

Different particle sizes and surface chemistries were chosen to investigate how root morphology and exudation is affected in various environments. The particle sizes chosen correspond to large soil particles (>2 mm), intermediary particles (53-250 µm), and small particles (<53 µm)(Six et al., 2000). Glass beads were chosen as they constitute an inert experimental system, for which the diameter of the spheres is defined, and the particles have a defined mineral composition (sizes: 3, 2, 1, 0.5 mm). The sand and clay substrates utilized here constitute more natural environments than glass beads (sand sizes: 4 mm, 250 µm, 5 µm; clay size: 4 mm). The sand and clay mineral composition was either defined by the manufacturer, or determined here. The sand substrates constituted of more than 98% quartz, whereas the clay was a mixture of 51% opal-CT, 37% mica-illite, 10% quartz, and K-feldspar and calcite traces (Fig S1).

To assess the chemical properties of the substrates, the sorption of metabolites to the substrates was assessed by incubating them with a defined medium, composed of more than forty metabolites belonging to various chemical classes also found in root exudates (amino acids, organic acids, sugars, bases/nucleotides/nucleosides, and others, see Table S2). The metabolite recovery rate from the substrates was determined by liquid chromatography/mass spectroscopy (LC/MS). The recovery rate from the glass beads and the 4 mm sand was comparable to the defined medium control without substrate, whereas the recovery rate from the 250 µm sand was approximately 30% lower, and from clay approximately 70% lower (Table S2, Fig. S7). Consistently, differences between clay and other substrates explained 84% of the variance in a principal component analysis, and only 8% of the variance accounts for differences between the control, glass beads, and sand (Fig. 1). The metabolites depleted by clay belonged to a variety of chemical classes, among them charged compounds such as organic acids and ammonium salts (carnitine, acetylcholine), other nitrogenous compounds (amino acids, nucleotides/nucleosides), as well as of comparatively polar compounds such as sugars (Table S2). We confirm that as expected, clay particles sorb a variety of metabolites from a defined medium.

**Figure 1.**
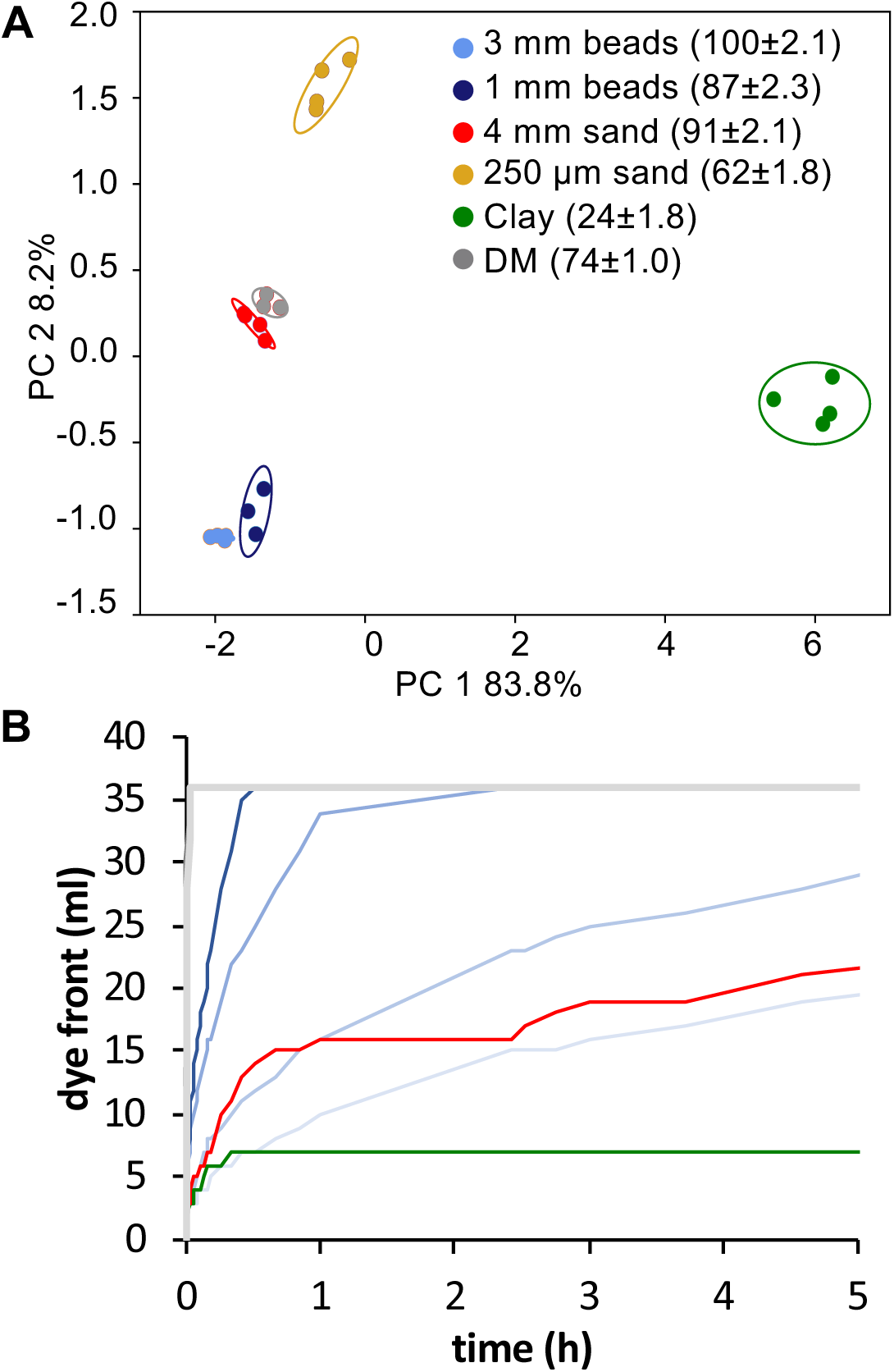
Physiochemical substrate properties (A) Principal component (PC) analysis of defined medium metabolites recovered from different substrates. The total metabolite recovery rate in percent is given for each substrate in brackets (means ± SEM, n=3). DM: defined medium control, no solid substrate. **(B)** Diffusion of colorimetric dye through various substrates as determined by the advancement of the dye front over time.

### Substrate diffusion rates

Altering particle size is expected to affect solute diffusion rates, and thus, may limit sorption. To investigate if exudation is limited by diffusion in our experimental system, diffusion rates of various substrates were determined using a colorimetric dye (Fig. 1). Diffusion was fastest in the controls without substrate added. The diffusion rate of the dye decreased with lower diameters in glass beads, whereas the diffusion rate in sand and clay was less linear. In 4 mm sand, the diffusion rate was initially similar to 1 mm glass beads, but then resembled more 0.5 mm beads. For clay, diffusion similar to 1 mm or 0.5 mm beads was observed initially, but subsequently, the dye front ceased moving, likely due to sorption of the negatively charged dye.

In our experimental setup, exudates required a minimum diffusion rate of 1.25 cm h^−1^ to reach the edge of the glass jar in which the plants were grown (10 cm jar diameter, 2 h exudate collection, plants positioned equidistant between center and edge). Thus, diffusion was not limiting in glass beads with a diameter equal to or greater than 1 mm, but might be limited in substrates with smaller diameters (Table S1).

This confirms, that as expected, sand and glass beads are inert substrates, whereas clay strongly sorbs a variety of metabolites. In addition, exudation may be limited by diffusion in substrates with particle sizes smaller than 1 mm.

### Root morphological changes in substrates

The aforementioned substrates with different particle size and chemistry were used to investigate how *Brachypodium distachyon* root morphology and exudation was affected in these conditions. Plants were grown for three weeks in the various substrates and in a hydroponic control. Exudates were collected, and root and shoot weight, as well as root morphology were assessed. Plants were grouped according to their behavior in the different substrates: plants with weights and root morphology similar to hydroponic controls were termed ‘big’ (big beads: 3 mm, 2 mm; big sand: 4 mm, 250 µm; clay), and plants with distinct weight and root morphology were termed ‘small’ (small beads: 1 mm, 0.5 mm, small sand: 5 µm. Fig. 2, grey areas).

**Figure 2.**
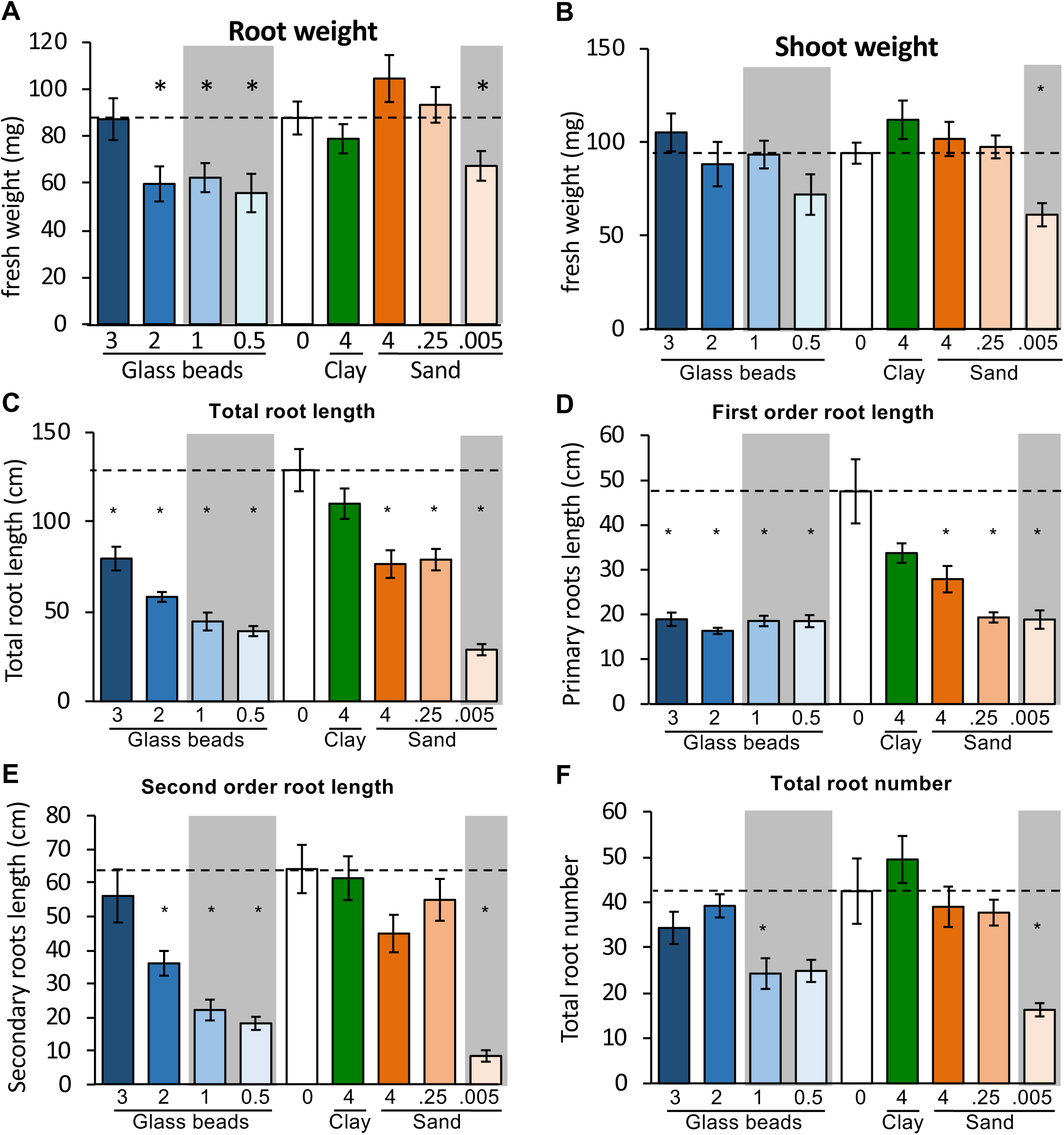
Tissue weight and root morphology. Root **(A)** and shoot fresh weight **(B)** of 3-week-old *B. distachyon* growing in different substrates. Root morphology was assessed as total root length **(C)**, which is the sum of first order root length **(D)**, second order root length **(E)**, and third order root length (Fig. S2); and the total root number **(F)**, which is the sum of first, second, and third order root numbers (Fig. S2). Particle sizes are indicated in cm. Data are means ± SEM, n>5. Significant differences are displayed as asterisks (*, ANOVA, p = 0.05) of substrates compared to hydroponic control (0, dashed line). Grey areas: plants with weight and root morphology distinct from hydroponic controls.

The root fresh weight of plants grown in 3 mm glass beads, 4 mm sand, 250 µm sand, and clay was comparable to the hydroponic control, whereas roots grown in 2 mm, 1 mm and 0.5 mm glass beads and 5 µm sand were significantly smaller (Anova/Tukey test, alpha = 0.05, Fig. 2). The shoot fresh weight of plants grown in 5 µm sand were significantly smaller compared to plants grown in all other conditions (Fig. 2). The altered root and shoot weights resulted in a decreased root/shoot ratios of 2 mm and 1 mm glass beads-grown plants, and an increased ratio of 5 µm sand-grown plants (Fig. S2).

Root length and number was assessed for first order roots (primary and crown roots), second order roots (laterals of primary and crown roots), and higher order roots. The total root length correlated with particle size, with maximal length for hydroponically and clay-grown roots, 30% shorter root systems for 3 mm beads-, 4 mm sand- and 250 µm sand-grown roots, and 50% or shorter root systems for 1 mm beads-, 0.5 mm beads-, and 5 µm sand-grown roots (Fig. 2). First order root lengths were significantly decreased by an average of 50% for all substrates except for clay (Fig. 2), whereas the second order root length was decreased by 40-70% in 2 mm beads-, 1 mm beads-, and 0.5 mm beads-grown roots, and by 85% in 5 µm sand-grown roots (Fig. 2). Higher order root lengths varied more within one experimental treatment, with a trend for higher total lengths in hydroponics and clay compared to glass beads and sand, and significantly lower lengths in 5 µm sand (Fig. S2).

Interestingly, root length had a higher Pearson correlation coefficient when correlated with particle size than root numbers. Only roots grown in 1 mm beads and 5 µm sand showed a statistically significant reduction in root number compared to hydroponic controls, which is a result of the large variability in total root number of hydroponically-grown plants (21 to 84 roots, Fig. 2, Fig. S3). The observed reduction in root number originated from a reduced number of secondary and higher order roots (Fig. 2, Fig. S2).

A correlation analysis between root and shoot weight, total root length, and total root number of all samples showed a significant correlation of all parameters investigated (Pearson correlation coefficients, p = 0.01, Fig. S4). Root weight and length, and to a lesser degree root number, correlated with particle size.

Overall, clay-grown plants were most similar to hydroponically-grown plants regarding tissue weight and root morphology. Plants grown in 3 mm glass beads or 4 mm sand had comparable fresh weight compared to the aforementioned plants, but slightly reduced total root length driven by a reduction in first order root length. Plants grown in 1 mm and 0.5 mm glass beads exhibited reduced root weight and root length, caused by a reduction in first and second order root length. Plants grown in 5 µm sand exhibited the largest reduction in tissue weight, root length and number.

### Spatially distinct exudation patterns

To learn more about how changes in root length and root number affect exudation, spatial patterns of exudation were investigated with Mass Spectrometry Imaging using Matrix Assisted Laser Desorption Ionization (MALDI) MS. The data was investigated manually, resulting in a total of 24 ions present in the vicinity of roots (Fig. 3, Fig. S5). There are some caveats to this analysis: A reliable identification of these ions is challenging, given that the MALDI is not well suited for fragmentation in the low-mass range and thus, no fragmentation patterns were acquired. In addition, MALDI data is subject to signal suppression and cannot easily be compared to LC/MS metabolite profiles due to the differing ability of the instruments to detect different ions. However, even without identifying the various ions, clear differences in spatial patterns could be observed. Some ions concentrated around root tip and elongation zone, supporting a role of these young root tissues in exudation. Some ions were detected along most of the root axis, suggesting exudation also from older root parts, whereas other ions were root-associated, which could either indicate short diffusion distances, or association with the cell wall. Overall, this data suggests that multiple tissues are involved in exudation.

**Figure 3.**
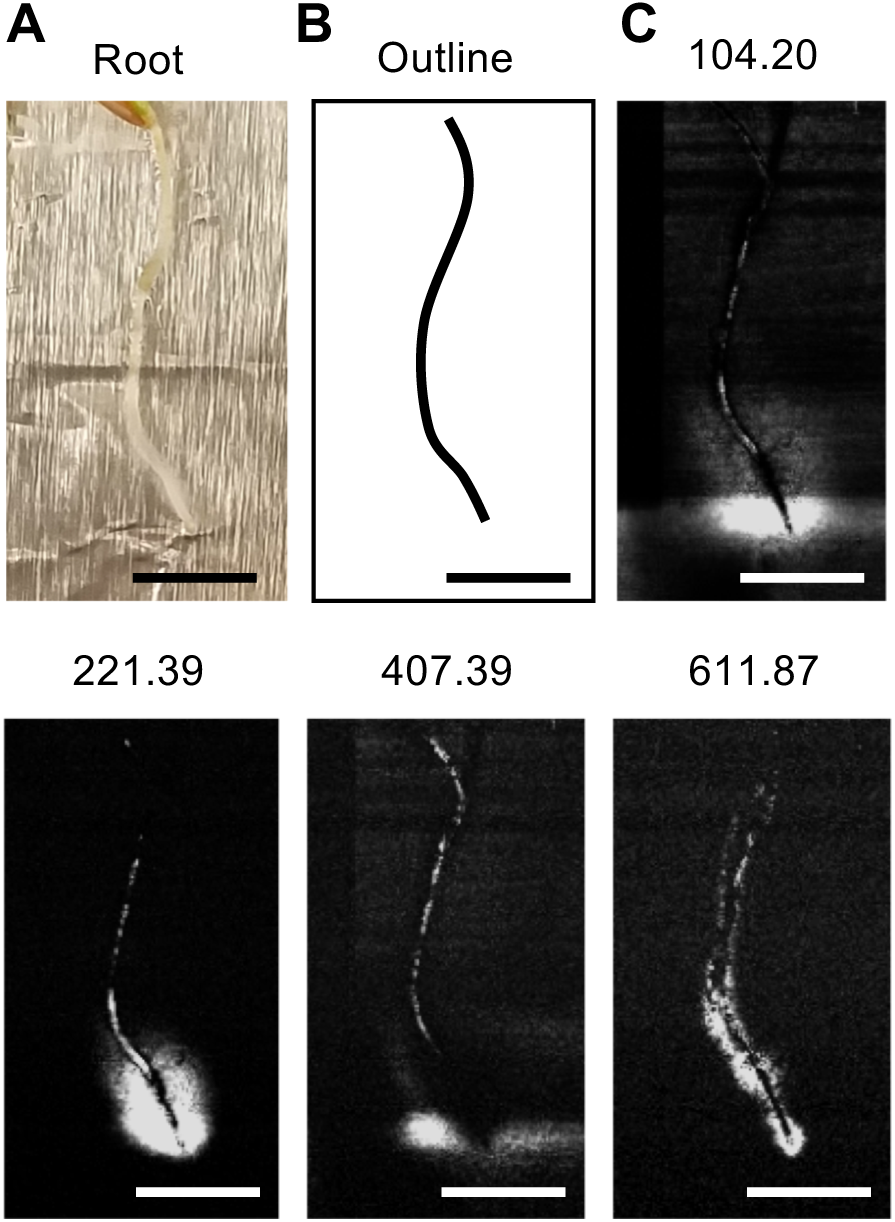
Spatial exudation patterns. Plants were incubated for 6 h to allow for exudation **(A)**. A traced outline of the root is displayed **(B)**. Distinct root-associated patterns of several ions were observed, the weight of the different ions observed is indicated above the panels **(C)**. The panels are a subset of Fig. S5. Scale bars: 1 cm.

### *In situ* vs. *in vitro* exudate profiles

Exudates were collected from plants grown in sterile conditions in the various substrates, to investigate whether the altered root morphology, and the various spatial exudation patterns resulted in an overall change in exudation profile. To be able to distinguish the effect of particle chemistry and altered plant metabolism, exudates were collected in two steps: first, exudates were collected from plants growing in the respective substrates (‘*in situ’*), and second, plants were removed from the substrate to collect exudates in a liquid medium (‘*in vitro’*). The first collection generated exudation profiles shaped by plant metabolism and particle chemistry, whereas the second collection generated exudate profiles shaped only by plant metabolism.

With both collection steps, approximately 100 metabolites were identified based on comparison of retention times, exact mass, and MSMS fragmentation patterns compared vs. authentic standards. The metabolites included organic acids, amino acids and derivatives, sugars and other carbohydrates, nucleic bases, nucleosides and nucleotides, among others. Multivariate statistical analysis revealed that 35% of *in situ* exuded metabolites were statistically significantly different in pairwise comparisons, compared to only 8% of *in vitro* exuded metabolites (Anova/Tukey test, p = 0.05, Fig. S7). A similar result was obtained when grouping the data in a hierarchical clustering analysis, with *in situ*-collected exudates clustering according to biological replicates, and *in vitro*-collected exudates clustering in a less pattern (Fig. S8).

Most of the variation for *in situ* exudates was explained by differences between exudates collected in clay, compared to other conditions, which is evident from a principal component analysis (Fig. 4). Similarly, in pairwise comparisons, clay-collected exudates showed most distinct metabolites (4-16), followed by 5 µm sand-collected exudates with 8-13 distinct metabolites (Fig. S7). In contrast to *in situ* exudates, *in vitro* exudates exhibited similar metabolite profiles when analyzed with a principal component analysis (Fig. 4, Fig. S8), and fewer than four metabolites had statistically significant abundances in pairwise comparisons (Fig. S7). Notably, *in vitro* exudates of clay-grown plants did not show more distinct metabolites in pairwise comparisons than plants grown in other substrates, suggesting that the *in situ* differences observed between exudate profiles of clay-grown plants and plants grown in other conditions resulted from the presence of the clay, and not from an altered plant metabolism.

**Figure 4.**
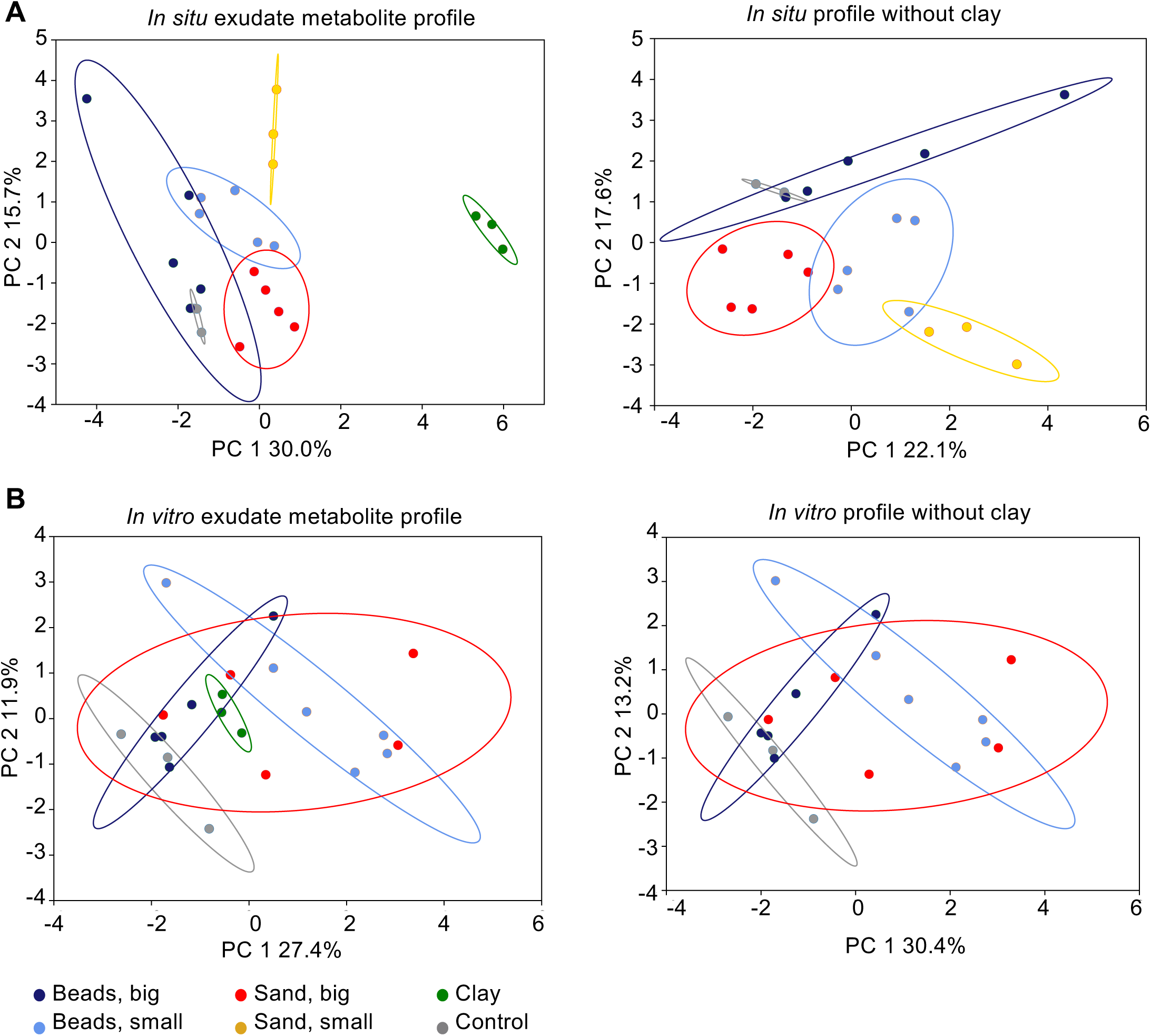
*In situ* and *in vitro* exudate metabolite profiles. Principal component (PC) analysis of root exudate metabolite profiles collected *in situ* **(A)** or *in vitro* **(B)**. Substrates are grouped as follows: beads big: 3 mm, 2 mm glass beads; beads small: 1 mm, 0.5 mm glass beads; sand big: 4 mm, 250 µm sand; sand small: 5 µm sand. The small sand dataset is missing from the *in vitro* analysis due to a technical error. Control: exudates from hydroponics. PC analyses of single substrates, and hierarchical clustering of the data are displayed in Fig. S8.

### *In situ* big vs. small sand exudate profiles

To further investigate differences of *in situ* collected exudates, metabolic profiles of the groups ‘big beads’, ‘small beads’, ‘big sand’, and ‘small sand’ were compared (grouping according to root morphology phenotypes)(Fig. 4, plots for individual substrates are shown Fig. S8). In a principal component analysis, exudate profiles of hydroponic and ‘big beads’ exudates overlapped, as well as ‘big beads’ and ‘small beads’. However, ‘big sand’ and ‘small sand’ grouped further apart (Fig. 4). Pairwise comparisons showed four distinct metabolites between ‘big beads’ and ‘small beads’, and nine distinct metabolites between ‘big sand’ and ‘small sand’. For the latter, all metabolites except thymine were more abundant in ‘small sand’. Among the more abundant metabolites were three phenolic acids (cinnamic, vanillic, 2,3-dihydroxybenzoic acid), a carboxylic acid (pyrrole-2-carboxylic acid), a sugar (mannosamine), a nucleotide (cytosine), and an amino acid derivative (carnithine)(Fig. S9). This data suggests that 5 µm sand has a higher retention rate for exudates than larger sand particles.

### *In situ* charged vs. uncharged substrates exudate profiles

To further examine the differences between clay-grown and hydroponically-grown *in situ* exudates, a multi variant test was used to compare metabolite abundances between the two conditions. Eighty-seven percent (20 of 23 metabolites) were significantly less abundant in clay than in hydroponics (Fig. 5). Most of these metabolites were nitrogenous, with two thirds containing a heterocyclic nitrogen group. Among these metabolites were nucleic bases / nucleosides / nucleotides with basic groups, amino acids with acidic or basic groups, a nitrogenous organic acid, and non-nitrogenous, aromatic organic acids. Three metabolites, two nitrogenous organic acids and a basic compound, were more abundant in clay-collected exudates (Fig. 5). These compounds were not detected in exudates of hydroponically-grown plants, in *in vitro*-collected exudates of clay-grown plants, or in clay control samples without plants (Table S2), which suggests that these compounds were released from clay only in presence of plants.

**Figure 5.**
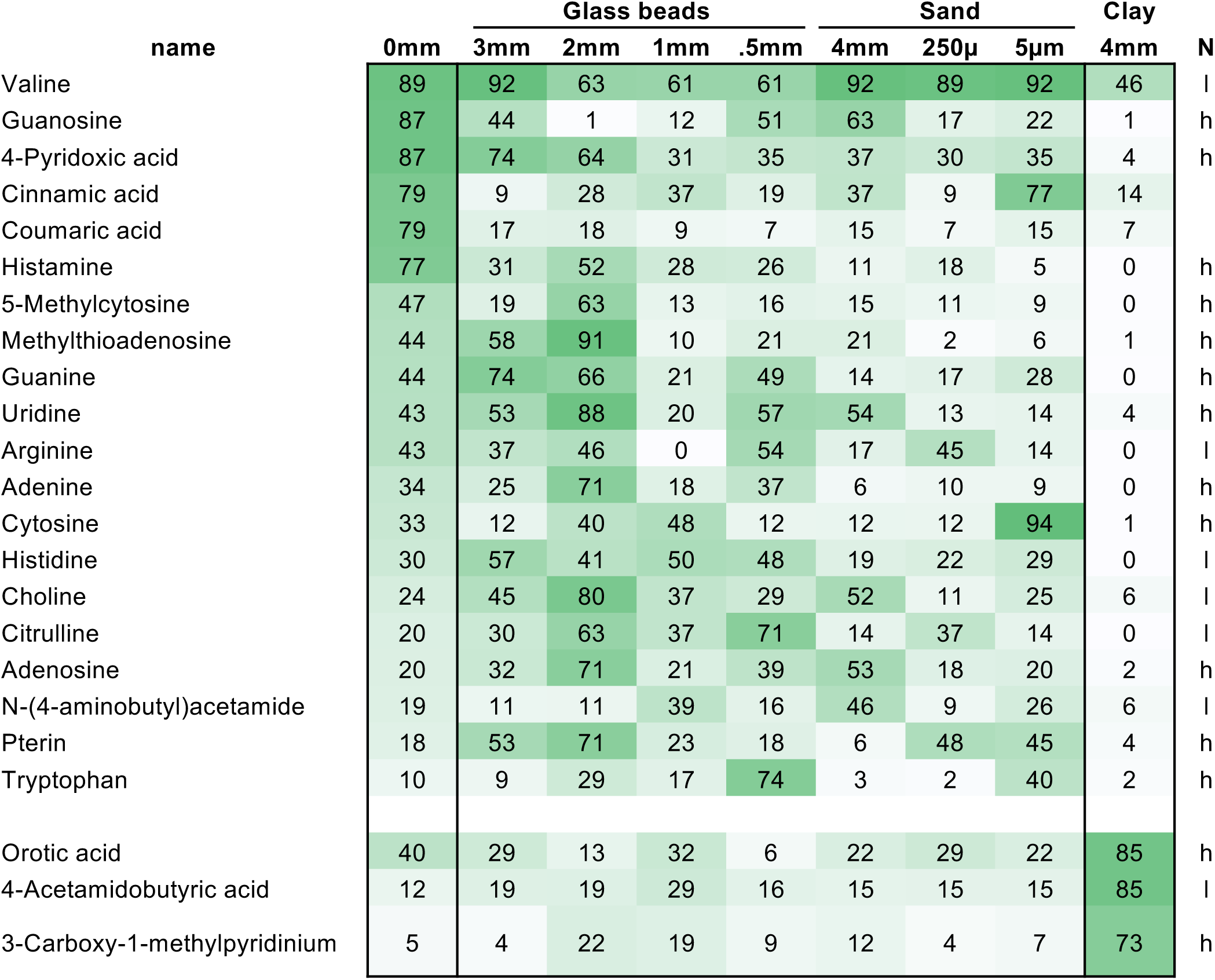
Distinct metabolites of *in situ* clay. Heatmap of normalized, averaged peak heights of metabolites significantly different between exudates collected *in situ* from clay-grown and hydroponically grown plants (black boxes). All substrates are displayed for comparison. Metabolites are sorted in descending abundance for hydroponic exudates (0 mm). Data are means of 3 biological replicates, ANOVA, p = 0.05. Nitrogenous compounds are labelled as containing heterocyclic (h) or linear (l) nitrogen. The full dataset is given in Table S1.

### A plant-beneficial microbe can grow on clay-sorbed metabolites

Since clay particles were found to strongly sorb exudate metabolites, we wondered if the sorbed metabolites were accessible to a plant-associated bacterium, supporting microbial growth. Thus, we first determined the desorption rate of metabolites from the various substrates by incubating glass beads (0.5-3 mm), sand (5 µm-4 mm), and clay (4 mm) with defined medium consisting of metabolites found in root exudates. The substrates were subsequently washed twice, and the recovered metabolites of all three steps were analyzed by LC/MS. As shown previously (Fig. 1), the metabolite recovery rate was comparable between the no substrate control and glass beads (63-80% recovery), lower in sand (31-51% recovery), and the lowest for clay (27%, Fig. S10). The metabolite recovery from washes was 4-14% for all substrates, suggesting that there are not more metabolites desorbing from clay than from other substrates.

Next, a rhizobacterium (*Pseudomonas fluorescens*) was incubated on various substrates pre-incubated with defined medium, and optical density (OD) was determined after three days. Defined medium with different concentrations was used as a positive control. Incubation of the rhizobacterium with 4 mm sand and 4 mm glass beads pre-incubated with defined medium resulted in the same OD as particles pre-incubated with water (Fig. S11), indicating that these substrates did not retain metabolites supporting growth. Incubation of the bacterium with clay pre-incubated with defined medium however did result in bacterial growth. As the control incubation of clay with defined medium (without bacteria) also showed a small increase in OD, presumably as a result of fine particles, the data presented in Fig. 6 is normalized by no-bacteria clay control samples (Fig. S11 presents the raw data).

**Figure 6.**
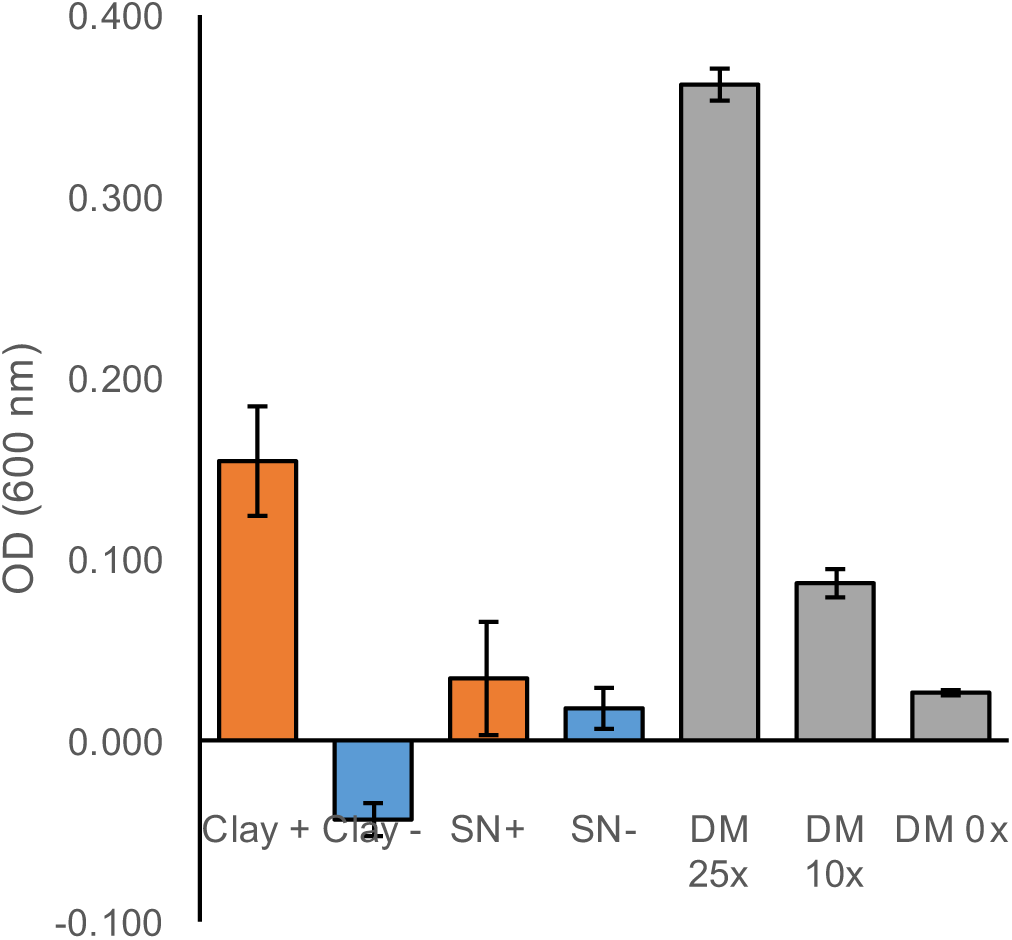
Rhizobacterium grows on clay-adsorbed metabolites. Optical density (OD at 600 nm) of *P. fluorescens* grown on clay pre-incubated with 50x defined medium (DM, Clay +) or 0x DM (Clay -), or in supernatant of clay pre-incubated with 50x DM (SN +) or 0x DM (SN -). All ODs are means ± SEM (n = 3), normalized by OD of clay without bacteria (see Fig. S11 for entire dataset). *P. fluorescens* growth in different concentrations of DM without substrate are given as comparison.

An additional control experiment consisted of incubating the defined medium-incubated clay with water for three days (the same timing as for the bacterial growth experiment), allowing for desorption of metabolites from clay particles. The supernatant was subsequently pipetted into a new well, bacteria were added, and allowed to grow for another three days. This experiment resulted in no bacterial growth (Fig. 6, Fig. S11), suggesting that bacterial presence is needed to desorb metabolites from clay particles. We conclude that this particular rhizobacterium is capable of desorbing exudate metabolites from clay to support growth.

## Discussion

### Root weight and morphology is influenced by substrate size

Growth of *B. distachyon* in particles with different sizes resulted in various morphological changes. A decrease in particle size resulted in decreased root weight, total root length, and in total root number, although the last parameter correlated less strongly (Fig. 2, Fig. S4). Root weight correlated positively with shoot weight, total root length, and total root number, indicating a dependency of the different parameters.

Notably, the morphology of *B. distachyon* grown in glass beads or sand were not directly comparable: plants grown in 5 µm sand had higher root weight and total root length than plants grown in 0.5 mm glass beads. Diffusion of a dye through glass beads and sand particles (Fig. 1) was lower in sand than in glass beads, suggesting that the sand particles might be arranged in a denser three-dimensional pattern compared to glass beads. The three-dimensional particle arrangement and other differences between the substrates, such as porosity, might account for root morphological differences observed between glass bead- and sand-grown plants.

For glass bead-grown plants, the reduction in total root length was explained by a reduction in second order root length, whereas the primary order root length was reduced in all sizes smaller than 3 mm. These trends for reduced root weight and root length but not root number are in line with observations made for maize grown in 1 mm vs hydroponic conditions (Veen, 1982; Boeuf-Tremblay et al., 1995; Groleau-Renaud et al., 1998). Interestingly however, whereas these studies noted an even larger decrease in shoot than in root weight, *B. distachyon* shoot weight did not change significantly in our experimental conditions. Similar results were found for lettuce grown in three different soils, where root fresh weight and morphology changed, but shoot weight was not affected (Neumann et al., 2014). The constant *B. distachyon* shoot weight might indicate sufficient nutrient uptake even by smaller root systems in the environments investigated. Thus, in future studies, it might be interesting to evaluate how the different root systems as generated here with different substrate sizes further respond to altered nutrient levels. One could expect an additional change in root morphology in different nutrient starvation conditions, as for example, root systems optimized for phosphate scavenging form numerous, short lateral roots, whereas roots optimized for nitrogen uptake exhibit fewer, but long lateral roots (Rellán-Álvarez et al., 2016). Phosphate movement is hindered by particles with a charged substrate, whereas nitrate movement is less affected by soil chemistry (Rellán-Álvarez et al., 2016). Thus, changes in root morphology of plants grown in phosphate-limited clay might be distinct from plants grown in phosphate-limited sand or glass beads. Plants grown in soil may exhibit additional changes in root morphology and metabolism, as shown for *B. distachyon* grown in a sterile soil extract, which showed reduced root length, elongated root hairs, and depleted a variety of metabolites from soil extract (Sasse et al., 2018). Root systems further respond to local alterations in soil structure, such as to the presence of micro- or macropores, or to air pockets (Rellán-Álvarez et al., 2016). Investigations of local root morphology responses in heterogenous settings with multiple, defined substrate sizes and chemistries will thus shed more light onto how plants respond to soil physiochemistry on a spatial and time scale. Multiple systems exist in which such experiments could be attempted, ranging from EcoFAB model systems to rhizotron designs (Grossmann et al., 2011; Parashar and Pandey, 2011; Rellán-Álvarez et al., 2015; Gao et al., 2018).

### Exudation across the root system

*B. distachyon* exhibited significantly altered root morphology when grown in particles with various sizes, with root weight and root lengths differing between conditions. The exudate profile however was indistinguishable for these plants when collected *in vitro* (Fig. 4). As the exudates were normalized by root fresh weight before measurement, this indicates that exudation per root fresh weight is constant. As root weight correlated with both, total root length and with total root number, it an additional method was needed to determine if the number of roots or the root length was important for exudation.

In the literature, root tips are often mentioned as predominant sites of exudation for several reasons: i) cell wall suberization of this young tissue is still low (Jones et al., 2009), ii)compounds, such as flavonoids, were imaged around root tips (Peters and Long, 1988), and iii) more microbes associate with tips compared to other root sections (DeAngelis et al., 2008). Few studies exist investigating spatial patterning of exudation, but some examples suggest that other tissues besides root tips might be involved in exudation: the localization of the malate transporter ALMT1 in Arabidopsis is confined to the root tip in untreated roots, but expands to the entire root system when treated with an activator, aluminum (Kobayashi et al., 2007), suggesting differential malate exudation from different parts of the root, depending on the environment. Strigolactone exudation is similarly environment dependent, with its transporter PDR1 expressed in single cells (hypodermal passage cells) along most of the roots (Kretzschmar et al., 2012). In addition, microbes do not only colonize root tips, but also prominently sites of lateral root emergence, and are found throughout the root system of plants (DeAngelis et al., 2008; Massalha et al., 2017). Also, distinct microbial populations are associated with *B. distachyon* seminal and nodal roots, as well as for nodal root tips vs nodal root bases(Kawasaki et al., 2016), which could be caused by differential exudation by these organs.

We used mass spectrometry imaging to investigate exudation across roots. This data cannot directly be compared to the root morphology and LC/MS data for technical reasons (see Methods) and the fact that the exudates were collected from three-week-old plants, whereas the imaging experiment was performed with seedlings due to space limitations. Some ions concentrated around the root tip, whereas others were also found in the root elongation and maturation zone, or all along the root axis. In addition, some ions were detected on the root itself, which could mean that they are part of the cell wall, or that they have a low diffusion speed. Further, the efficiency of detection for a specific compound differs between chemical imaging and liquid chromatography. Despite these limitations, our data suggest that root exudation is a spatially complex process. Exudation might take place in different ways: Root tip-exuded metabolites might diffuse, due to the absence of Casparian strips or secondary cell walls, or be transported. Metabolites exuded from older root tissues are more likely to be transported, either by channels facilitating diffusion, or by active transport proteins. It will be interesting in future studies to investigate the role of various root zones in exudation to determine which tissues are involved in exudation of various compounds. Further, it will be interesting to see if various root types produce different exudates.

### Root exudation is independent from particle size

Root exudate metabolite profiles were unaltered when plants were grown in different particle sizes. As the root weight, root number, and root length correlated, and the exudation of compounds was spatially complex, we conclude that exudation profiles are robust across different root morphologies. It might be interesting to investigate if exudation profiles change in plants with more radically altered root morphologies, in plants without secondary roots or root hairs, for example.

Exudation profiles were also comparable between plants grown in clay, sand, or glass beads, when collected *in vitro*. This suggests that the physiochemical environment does not alter plant metabolism, as long as other factors such as nutrient levels, light intensity, and humidity remain unchanged. However, the exudate metabolite profile of clay- vs. sand-grown plants was clearly different for *in situ* exudates (Fig. 4). A recent study found differences in sorghum exudates of plants grown in clay, sand, and soil (Miller et al., 2019). In this study, exudates were collected from roots with rhizosphere substrate still attached. The largest difference in this dataset was observed between soil-grown and sand- or clay-grown plants, which might be explained by soil-derived metabolites co-extracted with root exudates (Sasse et al., 2018; Miller et al., 2019). The authors showed some ions to be specifically up- or down-regulated in exudates of clay- vs. sand-grown plants, but their effect was not strong enough to separate the two conditions in a principal component analysis (Miller et al., 2019), which might be due to their exudate collection method, which was a mixture between the *in situ* and *in vitro* conditions utilized here.

Recently, it was suggested that root tips might detect the concentration of rhizosphere metabolites, altering root morphology and exudation accordingly (Canarini et al., 2019). Thus, clay-grown plants should exhibit an altered root morphology compared to hydroponically-grown plants, as clay sorbs a significant amount of exudates, changing the metabolite concentration around the root tip. However, the root morphology of clay-grown plants is statistically not different from hydroponically-grown plants (Fig. 2). However, only one particle size of clay was used here. We found that diffusion rates were limited in smaller particle sizes. In systems with low diffusion rates, exudate concentration is likely higher around the roots, which might lead to higher exudate re-uptake than in systems with larger particle sizes (Sherson et al., 2000; Sasse et al., 2018). It might thus be interesting to investigate if clays with different particle sizes might provoke a root morphology and exudation profile distinct from glass bead-grown plants. Substrate particle size might be a factor defining the amount of exudates present in soils.

### Exudates are strongly sorbed to clay

The largest difference in exudate profiles observed was between *in situ* clay-grown plants and other *in situ* conditions. Notably, the distinct exudation of clay-grown plants disappeared when exudates were collected *in vitro*, indicating that the differences observed resulted from the presence of clay, and not from an altered exudation of compounds by *B. distachyon*.

About 20% of compounds were distinct between hydroponic and clay exudates, and most of these compounds were reduced in abundance in the presence of clay. Among these compounds were organic acids, amino acids, nucleotides, and positively charged compounds. When clay was incubated with a defined medium, 75% of all compounds were reduced in abundance, among them negative and positive charged compounds, as well as neutral compounds. The higher metabolite retention by clay in the defined medium experiment compared to the plant experiment might be due to several factors: the clay was incubated for two hours with the defined medium, but for three weeks with plants producing exudates. Although exudates were also collected for two hours in the plant experiment, the clay was likely already saturated to some degree with exudates. The quantification of exudate amounts at different plant developmental stages in future studies would enable to estimate the total amount of compounds exuded, which would allow to correct for the difference in the two experimental setups.

The reduction of metabolite abundance in the presence of clay is most likely due to its high ion exchange capacity, compared to quartz-based particles such as sand or glass beads (Kabata-Pendias, 2004). Previous studies investigating sorption of bacterial lysates to ferrihydrite found a depletion of more than half of the metabolites (Swenson et al., 2015a). Similarly, incubation of bacterial lysates with a soil consisting of 51% sand, 28% silt, and 21% clay resulted in low metabolite recovery rates (Swenson et al., 2015b). These findings are consistent with our data.

Interestingly, three nitrogenous metabolites were higher in abundance in exudates of *in situ* clay-grown plants, two of which are organic acids, and a positively charged metabolite (orotic acid, 4-acetaminobutyric acid, 3-carboxy-1-methylpyridinium, Fig. 5). These compounds were not detected in clay negative controls, or in *in vitro* exudates of clay-grown plants, making it likely that the presence of plants lead to the release of these compounds from clay. Multiple examples exist in literature that describe a release of compounds from minerals by specific exudates. For example, plant-derived organic acids such as malate and citrate solubilize mineral-bound phosphate (Neumann and Martinoia, 2002), and plant-derived oxalate releases organic compounds bound to minerals, making them available to microbial metabolism (Keiluweit et al., 2015). Orotic acid has cytokinin-like activity (Shopova and Moskova-Simeonova, 2000), whereas the second compound is an acetylated form of gamma-aminobutyric acid, also with potential growth-regulating functions (Ramesh et al., 2016). The third compound is a pyridine derivative, which are precursors for secondary metabolite synthesis. The root morphology of plants grown in clay is comparable to hydroponically-grown plants (Fig. 2), which might indicate that the present concentration of these compounds is not sufficient to induce growth changes in clay-grown plants, or alternatively, that *B. distachyon* is not responsive to these signaling compounds. Altered exudation depending on the growth substrate was also described for tomato, cucumber, and sweet pepper growing in stonewool, with higher exuded levels of organic acids and sugars compared to glass bead-grown plants (Kamilova et al., 2006). The authors suggest that the presence of aluminium ions in stonewool might be responsible for the altered exudation observed. As the authors did not investigate *in vitro* collected exudates of stonewool-grown plants, it is unclear to which degree the observed effect was due to changes in plant metabolism, or due to the presence of stone wool.

In soils, metabolite sorption to minerals can lower decomposition rates (Baldock and Skjemstad, 2000). Also, the amount of clay in soil is correlated with retention of labeled carbon in soils (Baldock and Skjemstad, 2000). In clay-dominated soils, the size of clay particles shapes how much carbon can be retained: large clay aggregates were found to adsorb more carbon than smaller aggregates (Six and Paustian, 2014). Here, we only investigated one size of clay particles. It would thus be interesting in future studies to investigate the sorption behavior of clays with different particle sizes, and the ability of microbes to subsequently de-sorb these compounds. In natural systems, the presence of large amounts of clay with a specific particle size likely results in the sorption of plant-derived compounds to particles, changing the direct availability of these compounds to heterotroph organisms, and thus, altering soil processes.

### Microbial desorption of clay-bound metabolites

Microbes can release sorbed compounds from minerals, and they likely preferentially colonize minerals that are associated with compounds missing from the environment (Uroz et al., 2015). We tested if a plant-growth-promoting bacterium was able to grow on clay pre-incubated with a defined medium resembling root exudates. The rhizobacterium utilized in this study was indeed able to desorb metabolites from clay, utilizing them as a carbon source for growth (Fig. 6). In soils, root create zones with high metabolite density. The released exudates can either be directly taken up by root-associated microbes, or sorbed to minerals. The ability of root-associated microbes to release mineral-bound metabolites as an additional source for growth might be one trait that supports competitiveness and survival. Root-associated bacteria have distinct exudate substrate preferences from bulk soil bacteria (Zhalnina et al., 2018), which might also define the kind of compounds bacteria are able to release from minerals (Uroz et al., 2015). Our results are further evidence that minerals play an important role in plant-microbe interactions by sorbing root exudates, which can later be solubilized by microbes for growth.

We conclude that differently sized particles induce distinct root morphologies in *B. distachyon.* Root exudation was constant per root fresh weight, and the exudate metabolite profiles were robust across root morphologies. Mass spectrometry imaging showed presence of different ions across various regions of the root system, suggesting involvement of different tissues in exudation. Exudates were strongly sorbed by clay, significantly reducing the availability of free metabolites. Some of the clay-bound metabolites however could be utilized by a rhizobacterium for growth. Soil clay content thus is likely an important factor to consider when investigating root exudates or plant-microbe interactions in natural environments.

## Materials & Methods

### Substrates and particle sizes

Solid substrates used: Glass beads with sizes 3 mm, 2 mm, 1 mm, and 0.5 mm (Sigma-Aldrich 1040150500, 18406, Z250473, Z250465), sand with sizes 4 mm (Rock-It Direct LLC, 612 Silica Sand), 200-300 µm (‘250 µm sand’, Sigma-Aldrich SiO_2_274739), and 0.5-10 µm (‘5 µm sand’, Sigma-Aldrich SiO_2_ S5631), and clay with size 4 mm (Turface Athletics, MVP Calcined Clay). The manufacturer’s specifications for the clay are as follows: calcined, non-swelling illite clay with 60% minimum amorphous silica (SiO_2_), 5% Fe_2_O_3_, and less than 5% Al_2_O_3_, CaO, MgO, K_2_O, Na_2_O and TiO_2_; approximately 41% 2.38 mm (mesh 8), 16% 3.36 mm (mesh 6), 24% 1.68 mm (mesh 12), 18% 0.84 mm (mesh 20), and 1% <0.84 mm (mesh 30 and smaller); pH 6.5 ± 1.

All substrates were acid-washed with 0.1 M HCl for 1 h at 200 rpm, rinsed five times with MilliQ water, and baked for 30 min at 200°C.

### Sorption test with defined medium

The various substrates were sterilized, and incubated at 24°C for 8 hours with a 20 µm equimolar defined medium, encompassing a variety of chemical classes, among them amino acids, sugars, organic acids, and nucleotides (Jenkins et al., 2017). The sterility of the system was confirmed by plating an aliquot on LB plates, followed by a 3-day incubation. The defined medium was removed by pipetting, and the recovered volume recorded. Samples were filtered through a 0.45 µm filter (4654, PALL Life Sciences), and frozen at −80°C. See ‘Liquid chromatography mass spectrometry sample preparation’ for sample processing.

### Liquid chromatography mass spectrometry sample preparation

The frozen samples were lyophilized (Labconco FreeZone lyophilizer), resuspended in 3 ml LC/MS grade methanol (CAS 67-56-1, Honeywell Burdick & Jackson, Morristown, NJ), vortexed three times for 10 s, sonicated for 20 min in a water bath at 24°C, and incubated at 4°C for 16 h for salt precipitation. Samples were then centrifuged for 5 min at 5000 g, 4°C, supernatants were transferred to new microcentrifuge tubes, and evaporated at 24°C under vacuum until dry. Samples were resuspended in 500 µl LC/MS grade methanol, and the above procedure was repeated. Finally, samples were resuspended in 100% LC/MS grade methanol with 15 µM internal standards (767964, Sigma-Aldrich). The volume in the last step is relative to the amount of defined medium recovered (see ‘sorption test’), or relative to the root tissue fresh weight for root exudates (0.77 g ml^−1^).

### Diffusion experiment and exponential model

Pipettes with a 50 ml volume were sealed at the bottom with parafilm, placed vertically, 50 ml of substrate was added, and approximately 25 ml of 0.5x MS was added to immerse the substrate (there was no correlation between the amount of 0.5x MS added and the volume needed to immerse the substrate). Congo red 4B (C6277, Sigma-Aldrich) was solubilized in water at a concentration of 20 mg ml^−1^, and 250 µl of the dye was added simultaneously to pipettes containing the various substrates. The front of the dye was followed over time.

Fitting of an exponential model to the experimental data resulted in the determination of the exponential change factor (*b*), which describes the relative increase of effective diffusion coefficient (in cm^2^ h^−1^) and diffusion rate (in cm h^−1^), as 14.5 and 13.3, respectively. The initial value factor (*a*), which determines the asymptotic lower bound of the model at smaller bead sizes, was calculated as 0.5 for the effective diffusion coefficient, and as 0.388 for the diffusion rate.

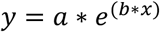

The experimental diffusion rates were determined by dividing the diffusion distance by the diffusion time. The effective diffusion coefficient was calculated for each substrate using the random walk diffusion coefficient equation below:

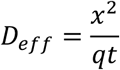

Where *x^2^* is the mean-square displacement, *q* is the dimensionality constant (set as 6 for 3-dimensional diffusion), and *t* is the time to maximum diffusion distance. This equation results in large effective diffusion coefficients for columns with large diffusion rates. The diffusion coefficient and the diffusion rate for 3 cm to 0.05 cm glass beads were modeled as:

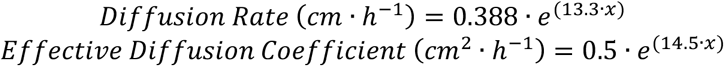

Where *x* is the glass bead diameter in cm, *e* is the base of the natural logarithm, 0.388 or 0.5 are the initial values of the modeled function for D or D_eff_ respectively, and 13.3 or 14.5 are the exponential change factors of the modeled function for D or D_eff_ respectively. The R^2^ value for these fits were 0.995 and 0.999 respectively, indicating a good fit of the data.

### Plant growth conditions

*Brachypodium distachyon* Bd21-3 seeds were dehusked and sterilized in 70% v/v ethanol for 30 s, and in 6% v/v NaOCl, 0.1% v/v Triton X-100 for 5 min, followed by five wash steps in water. Seedlings were germinated on 0.5x Murashige & Skoog plates (0.5x MS, MSP01, Caisson Laboratories; 6% w/v Bioworld Phytoagar, 401000721, Fisher Scientific, pH 5.7) in a 150 µmol m^−2^ s^−1^ 16 h light / 8 h dark regime at 24°C for three days.

Weck jars (743, Glashaus Inc.) were rinsed five times with MilliQ water, sprayed with 70% v/v ethanol, treated with UV for 1 h in a laminar flow hood, and dried over night. The jars were filled with 150 ml of the respective substrate, and 50 ml of 0.5x MS. Three seedlings were transferred into each jar, with the roots buried in the substrate. As a control, jars without substrate (0 mm, hydroponic control) were prepared: PTFE mesh (1100T45, McMaster Carr) was cut to fit the size of the jar, and autoclaved. Three openings were cut into the mesh to hold the seedlings. The mesh was transferred to jars with 50 ml 0.5x MS medium. For each condition, an experimental negative control was prepared containing substrate, but no seedlings. The experimental control jars were treated the same as the jars containing plants. To enable gas exchange, two strips of micropore tape (56222-182, VWR) were placed across the jar opening, the lid was set on top, and wrapped with micropore tape (56222-110, VWR) to ensure sterility. Plants were grown in a 16 h light / 8 h dark regime at 24°C with 150 µmol m^−2^ s^−1^ illumination, and the growth medium was replaced weekly: the old medium was removed by pipetting, and new 0.5x MS was added. Sterility of the jars was tested in week 3 by plating 50 µl medium on Luria-Bertani (LB) plates, following by three days incubation at 24°C.

### Root exudate collection

At 21 days after germination (21 dag), the medium in the jars was exchanged, and jars were incubated at 24°C with 150 µmol m^−2^ s^−1^ illumination for 2 h (‘*in situ’* treatment). Subsequently, plants were carefully removed from the substrate, and the roots of plants originating from the same jar were incubated submerged in 50 ml 0.5x MS for 2 h (‘*in vitro’* treatment) to account for the effect of the substrate presence on exudation (Fig. S1). Root exudates were passed through a 0.45 µm filter (4654, PALL Life Sciences), and frozen at −80°C. See ‘Liquid chromatography mass spectrometry sample preparation’ for sample processing.

### Root morphology imaging and analysis

After exudate collection, root systems were arranged on a glass plate, imaged, and root and shoot fresh weight was recorded. The root systems were traced with the Smartroot plugin (version 4.21) in ImageJ (version 2.0.0)(Lobet et al., 2011), and roots were assigned as first order (primary and crown roots), second order (lateral roots of primary and crown roots), and higher order (laterals of secondary roots). For each condition, a minimum of five root systems was analyzed.

Correlation analysis of root parameters were calculated using the excel pearson correlation coefficient (PCC) function, and the t-value was calculated using the following formula (n is the number of observations):

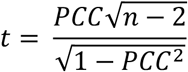

### MALDI Mass Spectrometry Imaging

Mass spectrometry imaging was used to determine if root exudate patterns varied across the root system. *Brachypodium distachyon* seeds were sterilized and germinated on 0.5 MS plates as described. A stainless steel MALDI plate was cleaned with 100% v/v ethanol, and a 7×7 cm square of aluminium foil was affixed to the plate with double-sided scotch tape. The foil was overlayed with 4 ml 0.1% ultrapure agarose (LS16500, Invitrogen, Toronto, ON, CA) to create a thin layer of agarose. Four-day-old seedlings were transferred to the agarose layer, and gentle pressure was applied with a spatula to embed the roots in the agarose. The stainless steel plate was transferred into a petri dish plate to keep humidity constant, and incubated for 6 h in a growth chamber with 150 µmol m^−2^ s^−1^ illumination and 24°C.

MALDI matrix was prepared as follows: 10 mg ml^−1^ a-cyano-4-hydroxycinnamic acid (70990, Sigma Aldrich, St. Louis, MO, USA) and 10 mg ml^−1^ Super-DHB (50862, Sigma Aldrich, St. Louis, MO, USA) were dissolved in 75% v/v methanol, 24.9% LCMS-grade water, 0.1% formic acid. The plate with the seedlings was removed from the growth chamber, leaves were cut to ensure flatness of the sample, and the sample was sprayed with MALDI matrix, which simultaneously desiccated the tissue. The plate was incubated for 24 h in a vacuum desiccator until completely dry. Mass Spectrometry Imaging was performed using a 5800 MALDI TOF/TOF (AbSciex, Foster City, CA, USA) in positive reflector MS mode with an Nd:YAG laser (200 Hz, 4400 laser intensity) acquiring spectra over a range of 50−2000 Da (1000 Da focus mass) and accumulating 20 shots/spot. The 4800 Imaging Tool software (Novartis and Applied Biosystems) was used to raster across the sample and record spectra in x-y step-sizes of 75 × 75 μm. Data viewing and image reconstruction were performed using OpenMSI (https://openmsi.nersc.gov) (Rübel et al., 2013).

### Liquid chromatography mass spectrometry methods and analysis

Metabolites were chromatographically separated with a hydrophilic liquid interaction chromatography on a Poroshell 120 HILIC-z 2.7 µm, 2.1 mm × 150 mm (Ag683775-924, Agilent Technologies) and detected with a Q Exactive Hybrid Quadrupole-Orbitrap Mass Spectrometer equipped (ThermoFisher Scientific). For chromatographic separations, an Agilent 1290 series HPLC system was used with a column temperature of 40°C, 3 µl sample injections, and 4°C sample storage. A gradient of mobile phase A (5 mM ammonium acetate, 0.2% acetic acid, 5 µM methylene-di-phosphonic acid in water) and B (5 mM ammonium acetate, 0.2% acetic acid, 95% v/v acetonitrile in water) was used for metabolite retention and elution as follows: column equilibration at 0.45 ml min^−1^ in 100% B for 1.0 min, a linear gradient at 0.45 ml min^−1^ to 11% A over 10 minutes, a linear gradient to 30% A over 4.75 min, a linear gradient to 80% A over 0.5 min, hold at 0.450 ml min^−1^ and 80% A for 2.25 min followed by a linear gradient to 100% B over 0.1 min and re-equilibration for an additional 2.4 min. Each sample was injected twice: once for analysis in positive ion mode and once for analysis in negative ion mode. The mass spectrometer source was set with a sheath gas flow of 55, aux gas flow of 20 and sweep gas flow of 2 (arbitrary units), spray voltage of |±3| kV, and capillary temperature of 400°C. Ion detection was performed using the Q Exactive’s data dependent MS2 Top2 method, with the two highest abundance precursory ions (1.0 m/z isolation window, 17,500 resolution, 1e5 AGC target, stepped normalized collisions energies of 10, 20 and 40 eV) selected from a full MS pre-scan (70-1050 m/z, 70,000 resolution, 3e6 AGC target, 100 ms maximum ion transmission) with dd settings at 1e3 minimum AGC target, charges excluded above |8| and a 7 s dynamic exclusion window. Internal and external standards were included for quality control purposes, with blank injections between every unique sample, and samples were injected in randomized order. Due to an instrumentation error, the *in vitro* exudate profiles of 5 µm sand-grown plants are missing.

### Metabolite identification and statistical analysis

Ion chromatograms corresponding to metabolites represented within our in-house standard library were extracted from LC/MS data with Metabolite Atlas (https://github.com/biorack/metatlas) (Bowen and Northen, 2010; Yao et al., 2015). Metabolites were identified following the Metabolomics Standards Initiative conventions, using the highest confidence level (‘level 1’, MSI identifications), which is identified as at least two orthogonal measures vs. authentic chemical standards (Sumner et al., 2007). In all cases three orthogonal measures were used: retention time (within 1 minute vs. standard), fragmentation spectra (manual inspection), and accurate mass (within 20 ppm). In general, accurate masses were within 5 ppm, though the error was higher for low mass ions in negative mode (but still below 20 ppm). Peak height and retention time consistency for the LC/MS run was ascertained by analyzing quality control samples that were included at the beginning, during, and at the end of the run. Internal standards were used to assess sample-to-sample consistency for peak area and retention times. Metabolites that did not show a signal three times higher in one experimental treatment compared to the experimental blanks were excluded. Metabolite peak heights were normalized by setting the maximum peak height detected in any sample to 100%, and expressing the metabolite intensity relative to this maximum peak height. This allows for relative comparison of peak heights between samples (e.g. if a compound of interest is significantly different between samples), but not for absolute metabolite level quantification (e.g. µg of a compound of interest per gram tissue). Chemical classes were assigned to metabolites with the ClassyFire compound classification system (Djoumbou Feunang et al., 2016).

To explore the variation between experimental conditions, the metabolite profiles were PCA-ordinated, and the 95% confidence level was displayed as ellipses for each treatment. Hierarchical clustering analysis with a Bray Curtis Dissimilarity Matrix was performed with the python 2.7 Seaborn package. Metabolite significance levels were analyzed with the python SciPy ANOVA test coupled to a python Tukey’s honestly significant difference test with alpha = 0.05 corresponding to a 95% confidence level.

### Rhizobacterium growth experiment

A selection of different substrates (3 mm glass beads, 4 mm sand, 4 mm clay) was incubated with 50 times concentrated defined medium (50x DM, see: “Sorption test with defined medium” section, (Jenkins et al., 2017)) or with 0x DM (DM without carbon sources, but with vitamins and minerals) for 6 h at 23°C. The substrates were subsequently washed three times with water, to remove soluble metabolites.

Substrates were added to a 12-well plate (2 cm^3^ each), and 2 ml of 0x DM was added to each well. A *Pseudomonas fluorescens* WCS415 preculture was grown in 5 ml 20x DM for 16 h at 30°C, 200 rpm. The culture was pelleted at 4000 g, 23°C for 5 min, and resuspended in 0x DM. The wells were inoculated with an initial optical density (determined at 600 nm) of 0.05 in triplicates. The plates were incubated at 30°C for 3 d (no shaking), 1 ml of the supernatant was removed to determine OD at 600 nm. Positive growth controls were *P. fluorescens* lines grown in the same experimental setup in 50x, 20x, 10x, and 0x DM, but without substrate. A set of negative controls was prepared to account for different variables in the experiment: Substrates incubated with 0x DM with bacteria were set up as a growth control, accounting for metabolites already adsorbed to clay. Substrates incubated with 50x and 0x DM but without bacteria were used to control for changes in optical density of clay caused by DM.

The metabolite desorption experiment was performed by adding 2 cm^3^ of clay pre-incubated with 50x or 0x DM to a 12-well plate in triplicates, followed by the addition of 2 ml of 0x DM. The plate was incubated for 3 d at 30°C. Subsequently, 1.5 ml of the supernatant was removed, and placed in a new 12-well plate. Half of the wells were inoculated with *P. fluorescens,* the other half served as negative controls. OD at 600 nm was determined after 3 d of growth at 30°C

---

**Figure S1.**
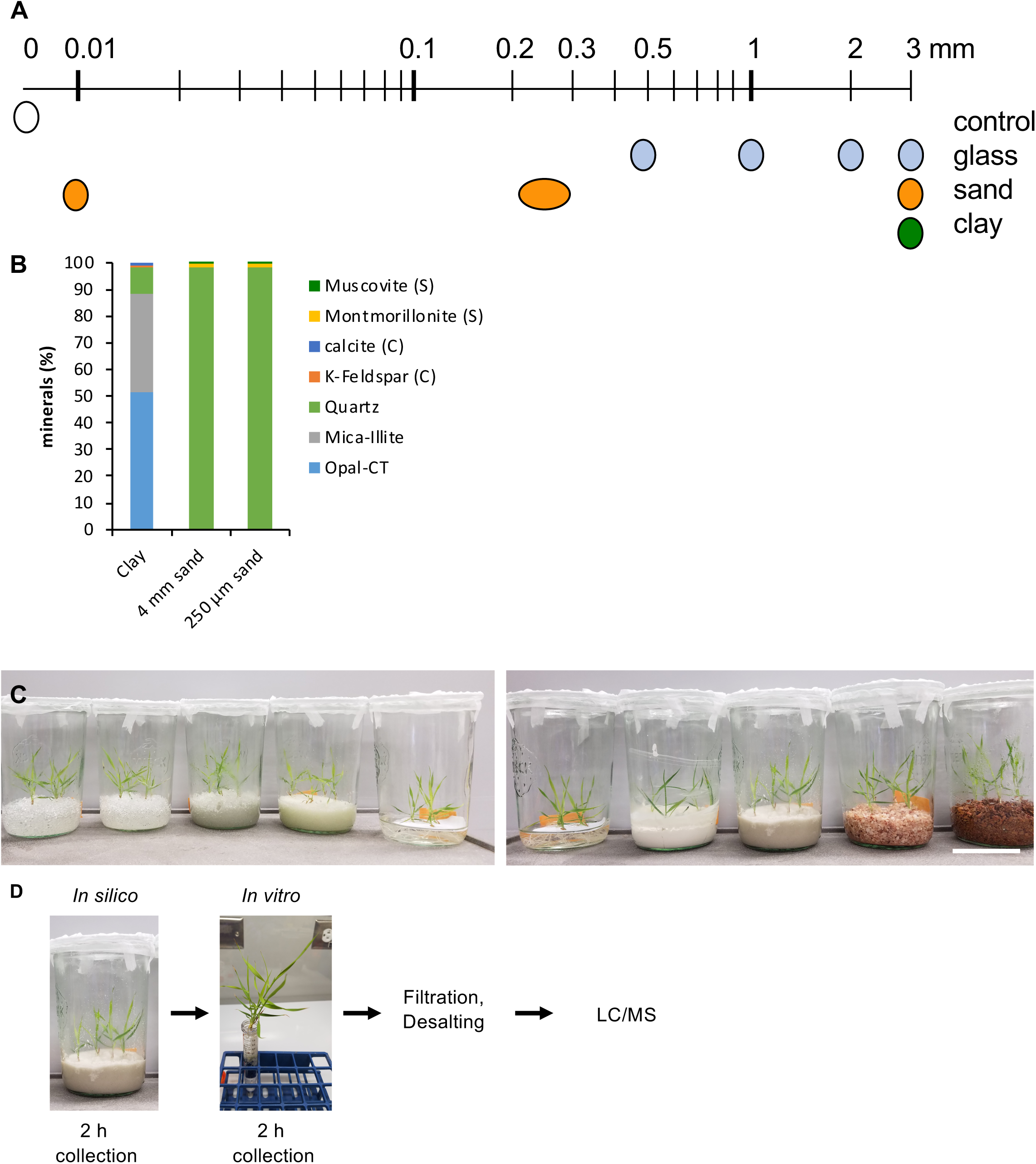
Experimental setup (A) Illustration of particle sizes utilized, logarithmic scale. **(B)** X-ray diffraction analysis of 4 mm clay, 4 mm sand, and 250 µm sand. Trace elements are indicated as being present in sand (S) or clay (C). **(C)** Jar setup, one jar per condition is depicted. Top panel, from left: 3 mm, 2 mm, 1 mm, 0.5 mm glass beads, hydroponic control. Bottom panel, from left: hydroponic control, 5 µm sand, 250 µm sand, 4 mm sand, 4 mm clay. Scale bar: 10 cm. **(D)** Root exudate collection procedure, the in vitro picture displayed is representative of the setup, but a larger tube (50 ml) was used for exudate collection.

**Figure S2.**
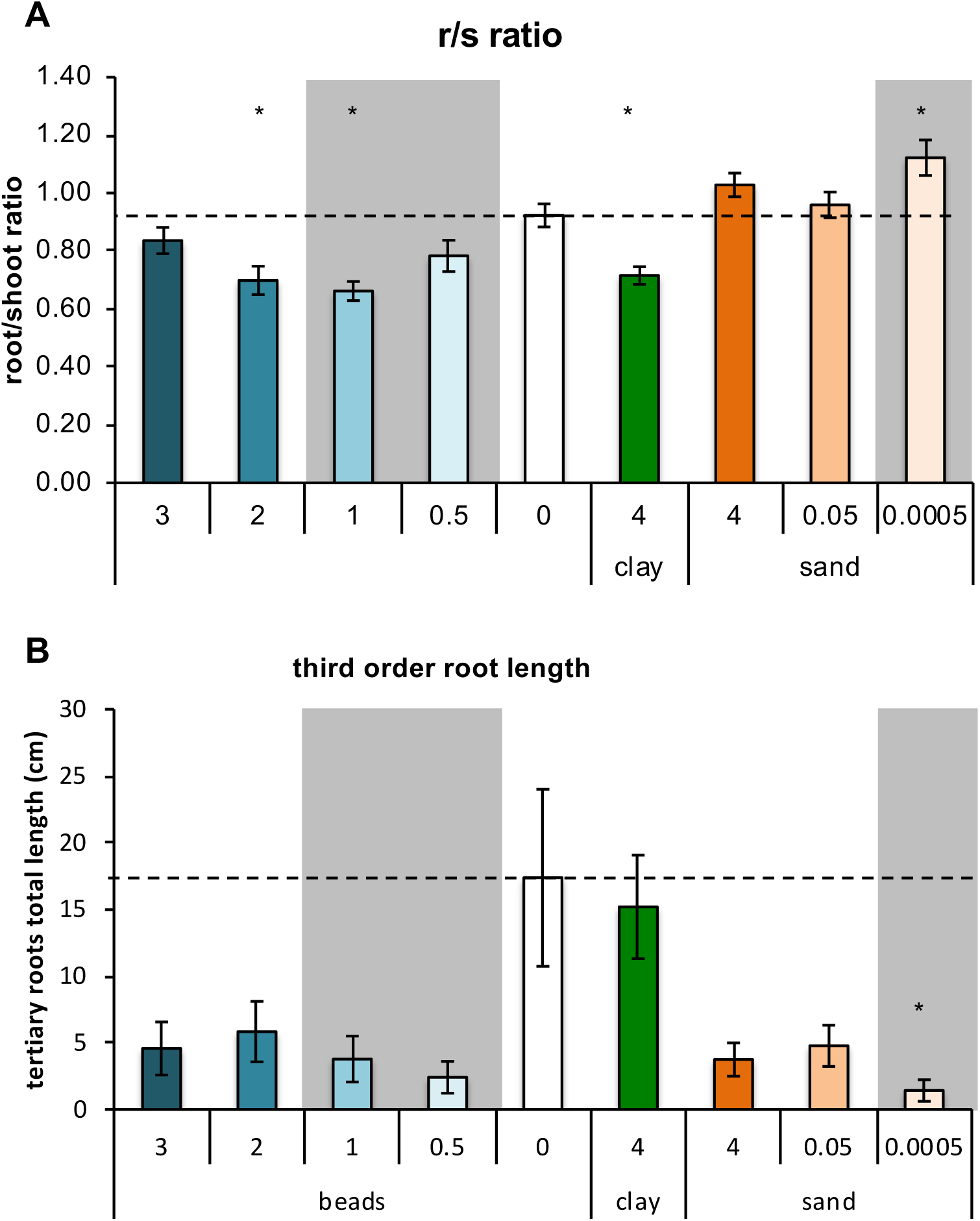
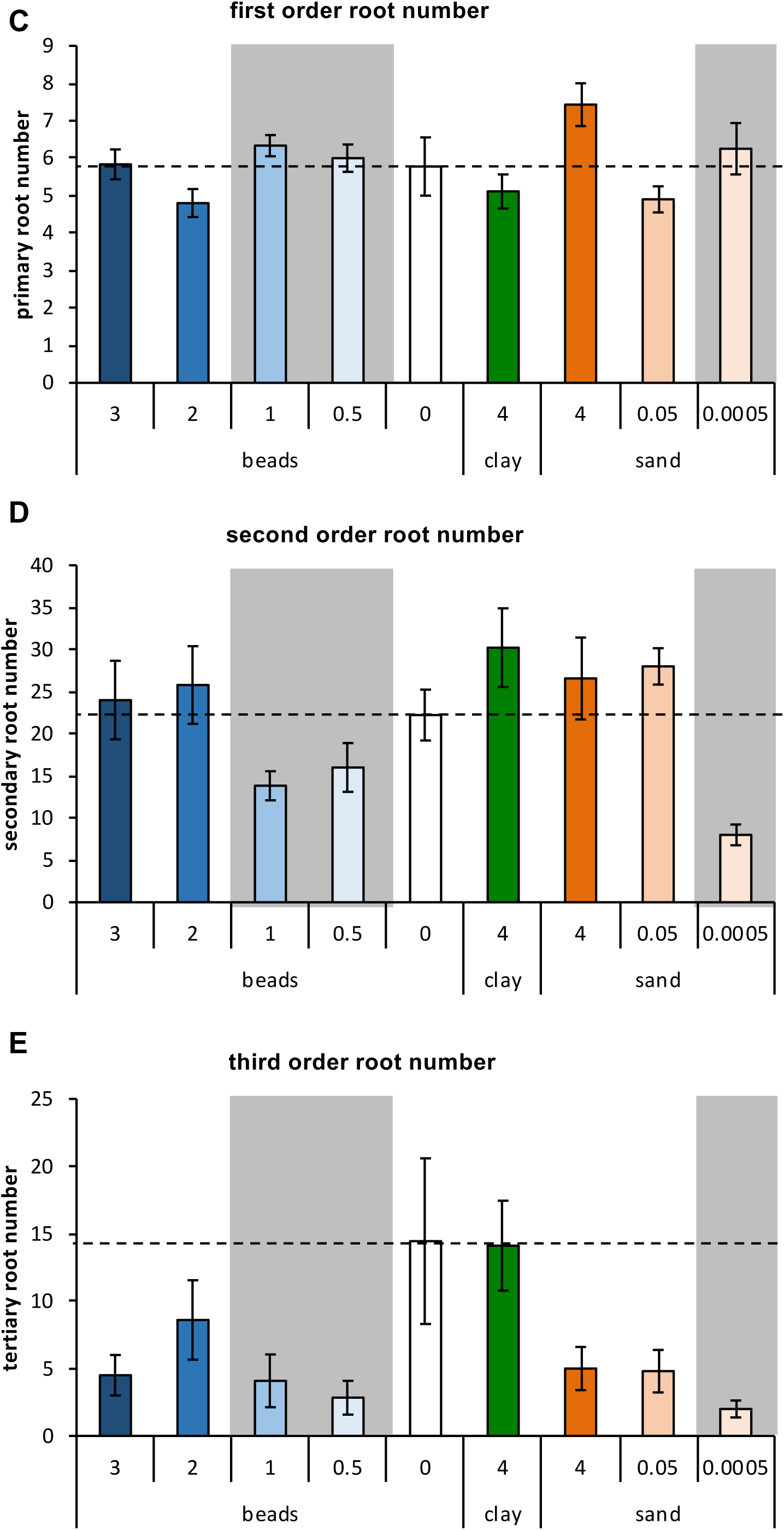
Additional root morphology. Root/shoot ratio **(A)**, third order root length **(B)** first order **(C)**, second order **(D)**, and third order **(E)** root numbers. Data are means ± SEM, n>5. Significant differences are displayed as asterisks (*, p = 0.05) of substrates compared to hydroponic control (0, dashed line). Tissue weights, total root length and number, first and second order root lengths are given in Fig. 2. Grey areas: plants with weight and root morphology distinct from hydroponic controls.

**Figure S3.**
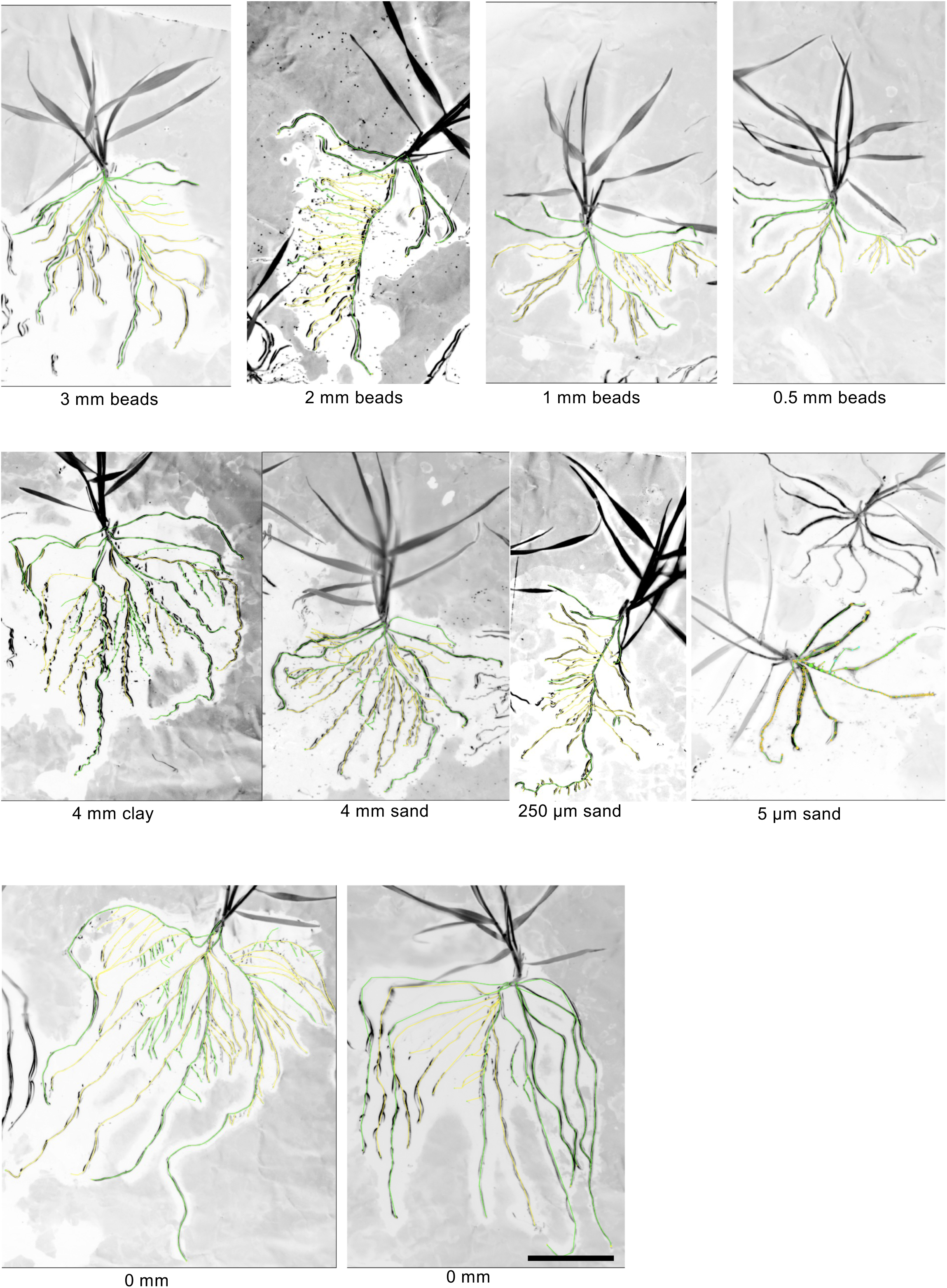
Representative root morphologies. Representative root morphologies of 3-week-old *B. distachyon* grown in the various substrates. First order roots were traced in green, second order roots in yellow, and third order roots in green (SmartRoot, (Lobet et al., 2011)). Quantification of root systems are displayed in Fig. 2 and Fig. S2. Scale bar: 3 cm.

**Figure S4.**
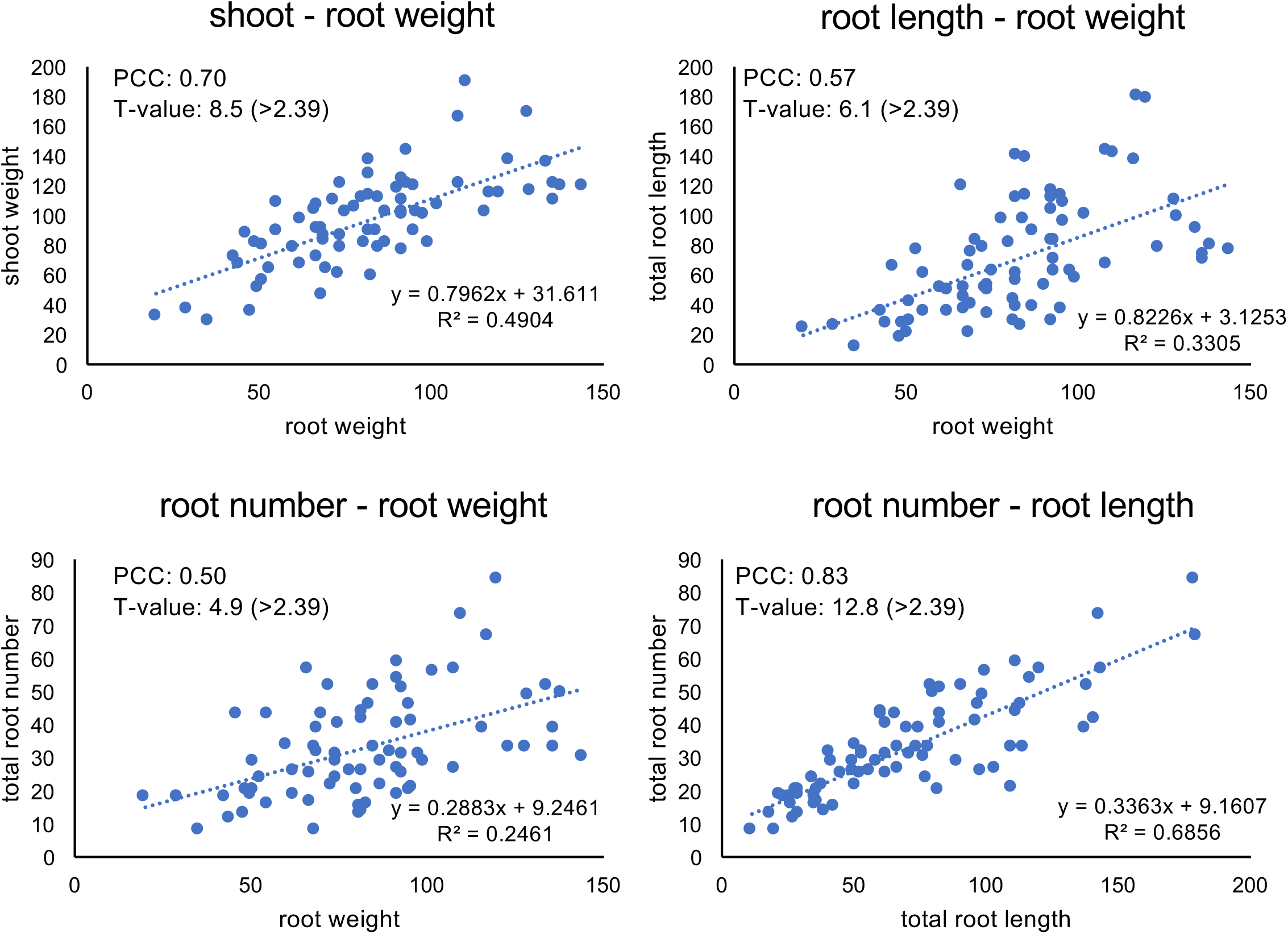
Correlation analysis of root parameters. Correlation plots of various root parameters. Dashed lines are linear regressions, equations and R^2^ values are given. Pearson correlation coefficients (PCC) were calculated, and the respective t-values are given.

**Figure S5.**
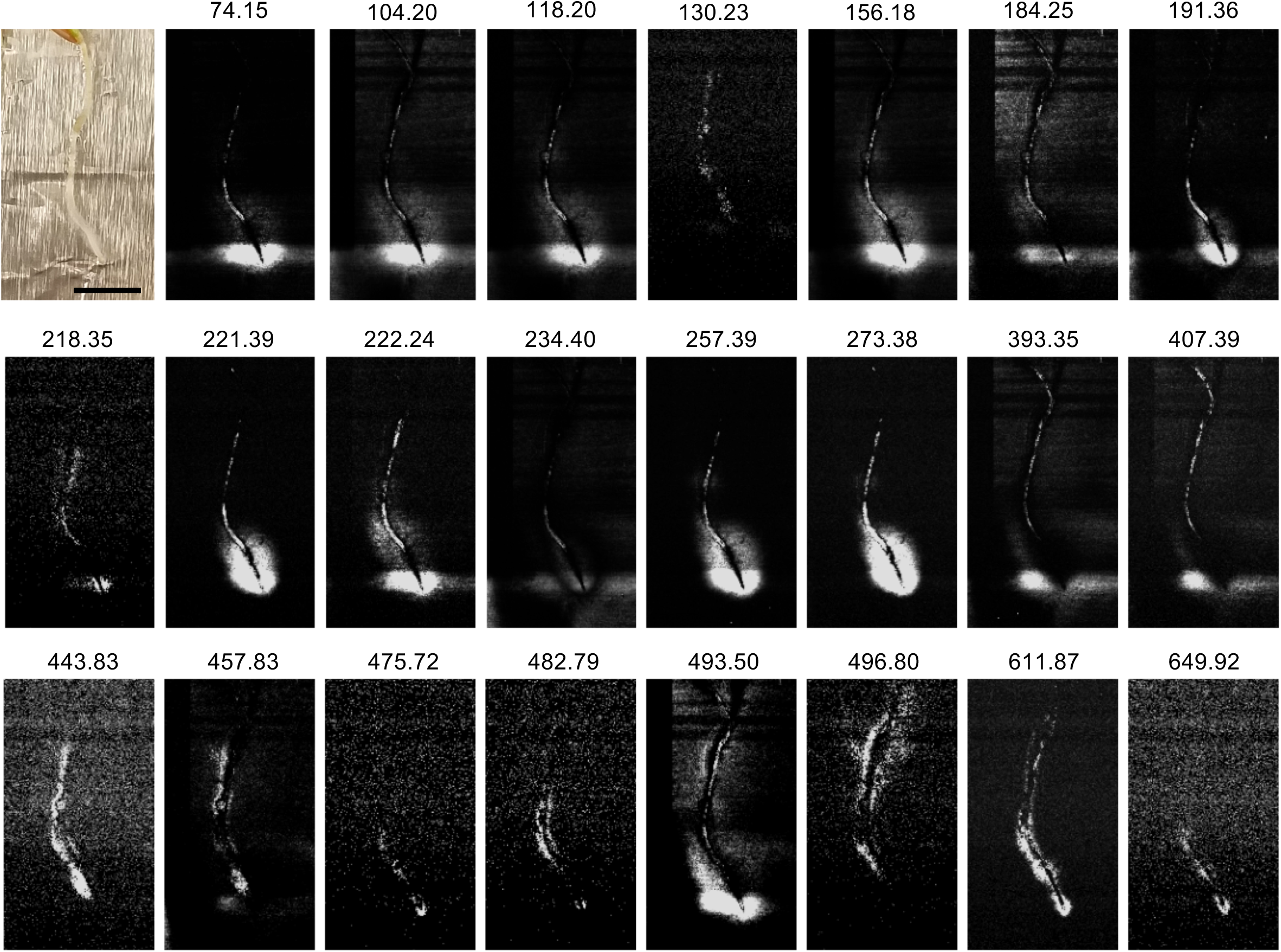
Additional spatial exudation patterns. Plants were incubated for 6 h to allow for exudation. Distinct root-associated patterns of several ions were observed, the weight of the different ions observed is indicated above the panels. Selected panels are presented in Fig. 3. Ion 156.18 is possibly a M+K-H adduct of 118.20. Scale bar: 1 cm.

**Figure S6.**
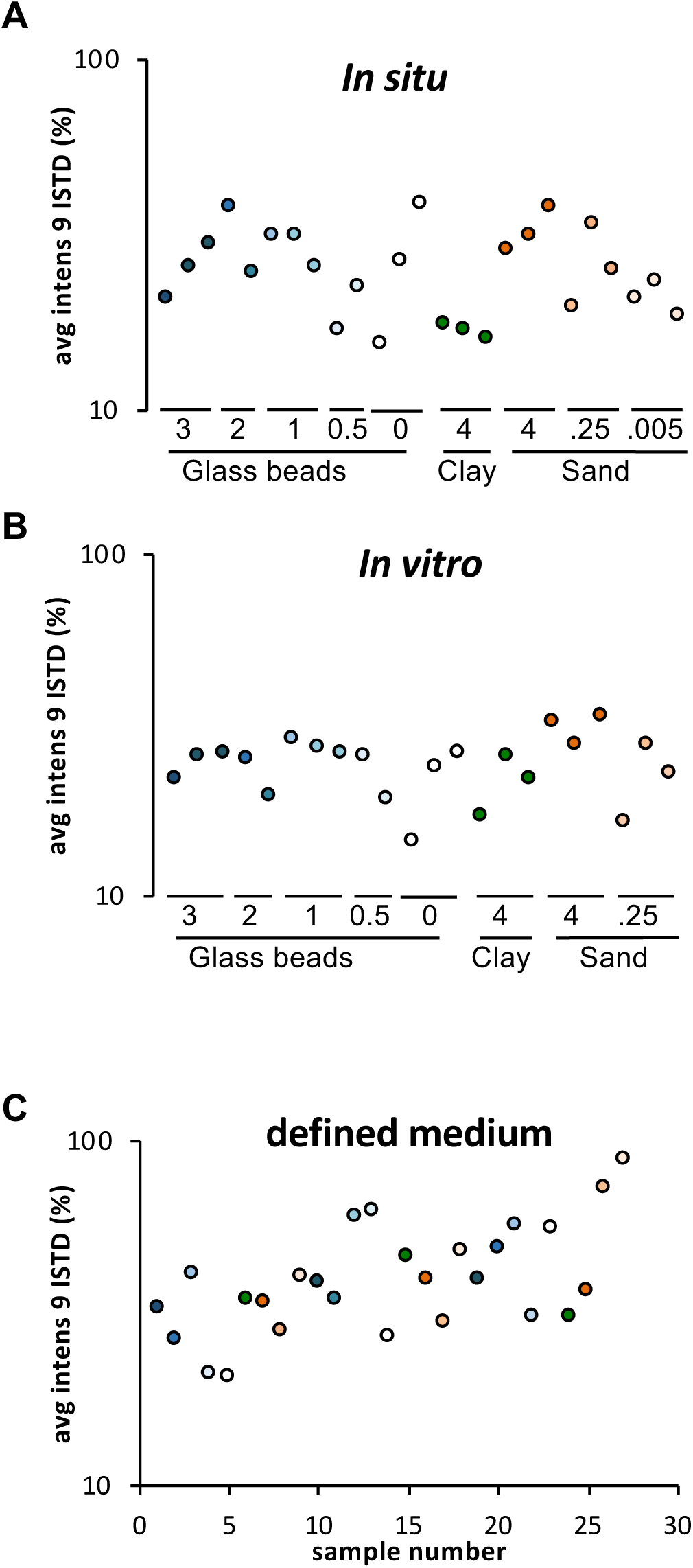
LC/MS internal standards. Average intensity of nine internal standards used for quality control for *in situ* collected exudates **(A)**, *in vitro* collected exudates **(B)**, and substrates incubated with defined medium **(C)**. Particle sizes are indicated in cm.

**Figure S7.**
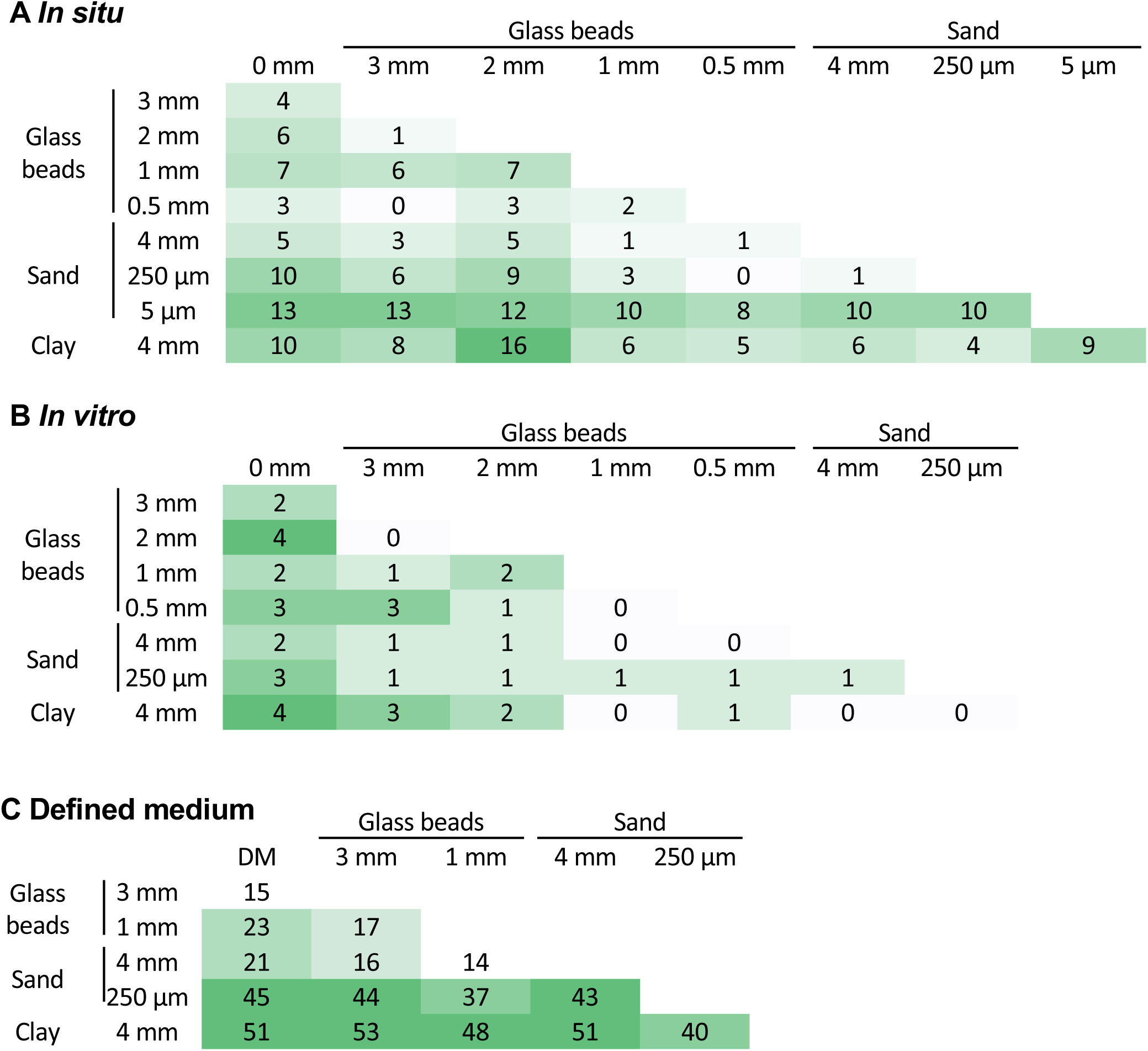
ANOVA-Tukey test results. Number of significantly different metabolites in pairwise comparisons of *in situ* collected exudates **(A)**, *in vitro* collected exudates **(B)**, and substrates incubated with defined medium **(C)**. Data are means of 3 biological replicates, ANOVA, p = 0.05.

**Figure S8.**
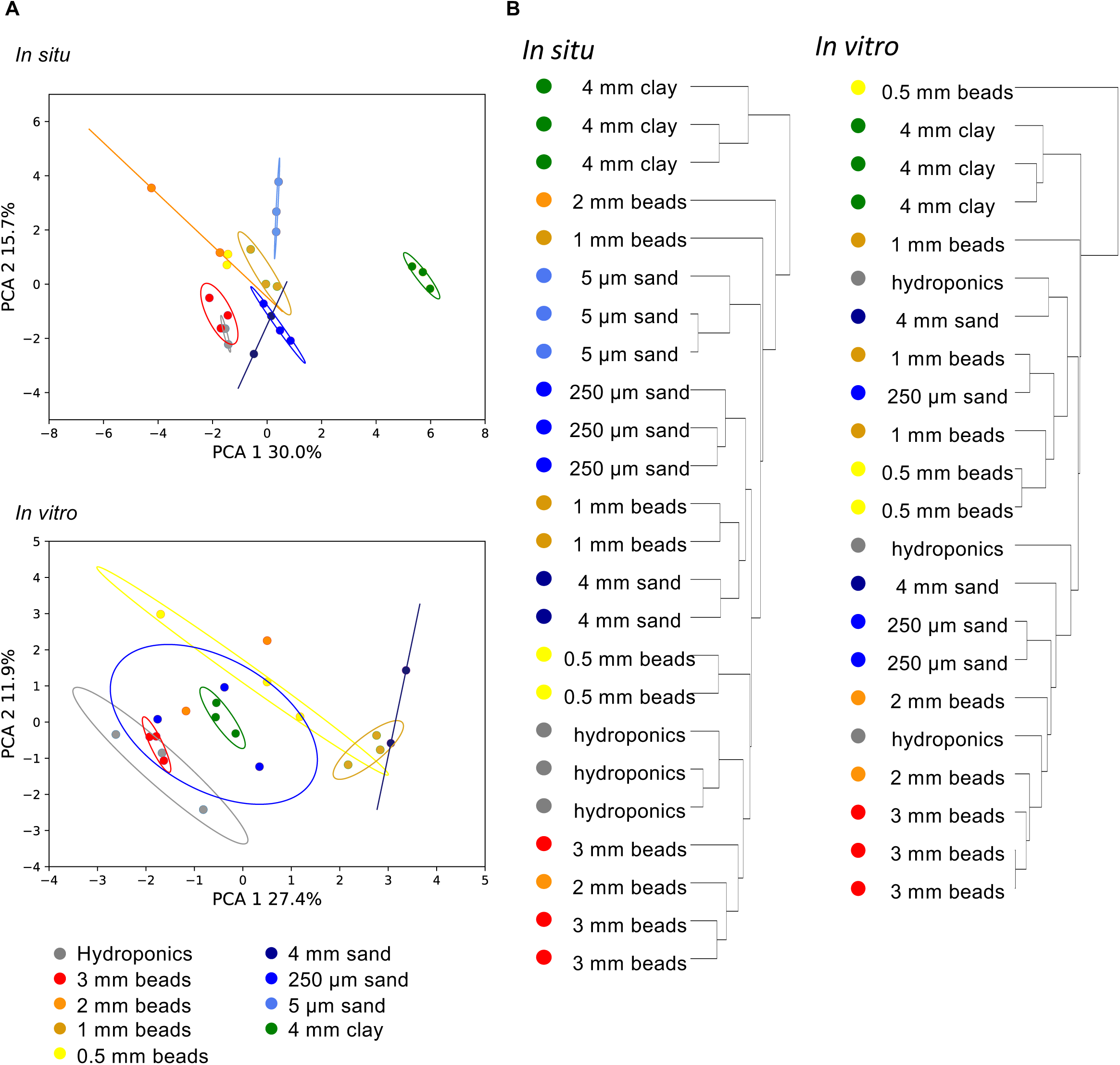
*In situ* and *in vitro* exudate metabolite profile analyses. Principal component (PC) analysis of metabolite profiles of root exudates collected *in situ* or *in vitro* **(A)**, and hierarchical clustering of the same datasets **(B)**. PC analyses of the data grouped by root morphology is displayed in Fig. 4.

**Figure S9.**
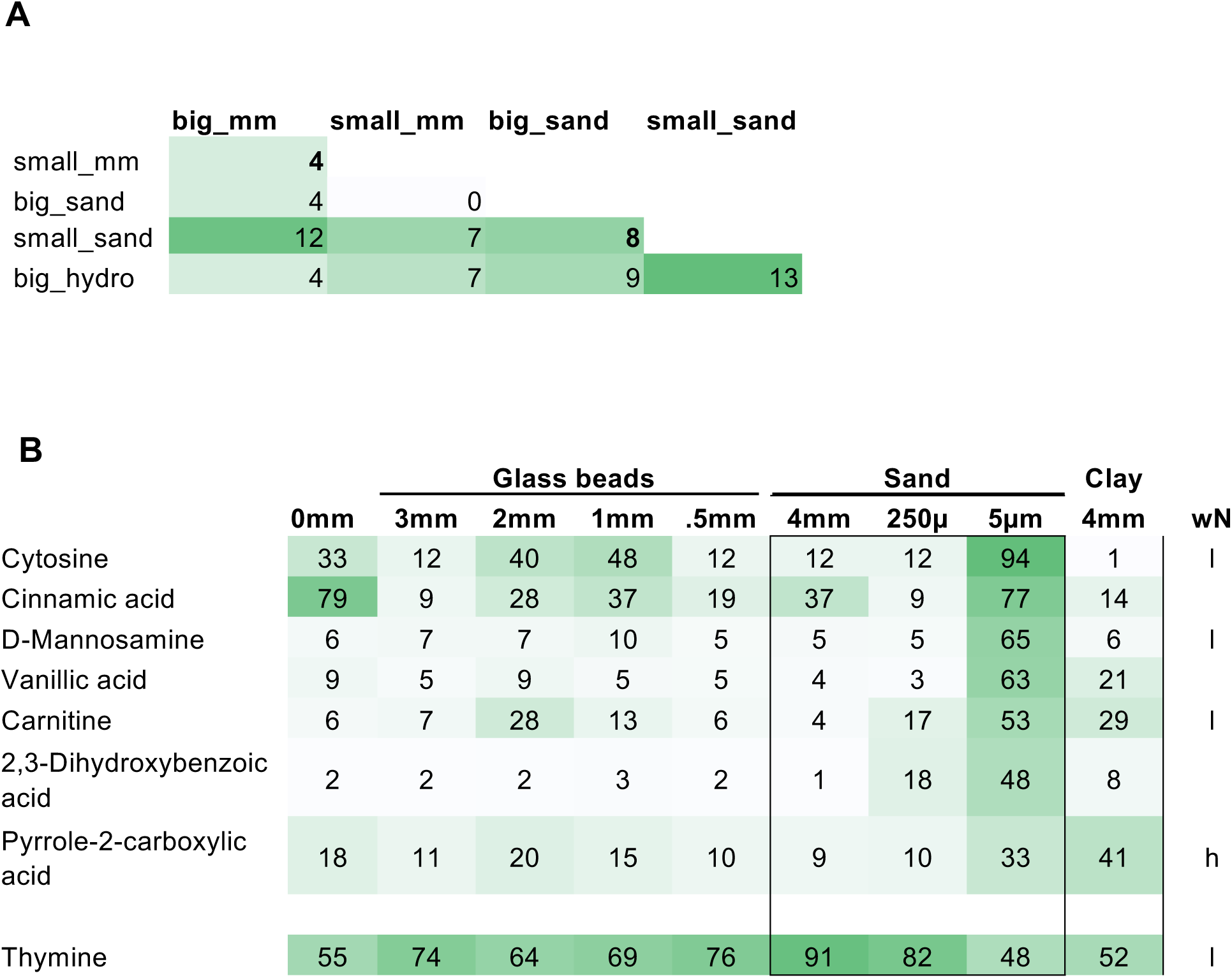
Distinct metabolites in big vs small sand (A) Number of significantly different metabolites in pairwise comparisons of metabolite profiles grouped by root morphology (ANOVA, p = 0.05). Substrates are grouped as follows: beads big: 3 mm, 2 mm glass beads; beads small: 1 mm, 0.5 mm glass beads; sand big: 4 mm, 250 µm sand; sand small: 5 µm sand. **(B)** Significantly different metabolites in 5 µm sand vs 4 mm sand (ANOVA, p = 0.05). The other datasets are displayed for comparison. Nitrogenous compounds are labelled as containing heterocyclic (h) or linear (l) nitrogen. Full dataset is given in Table S1.

**Figure S10.**
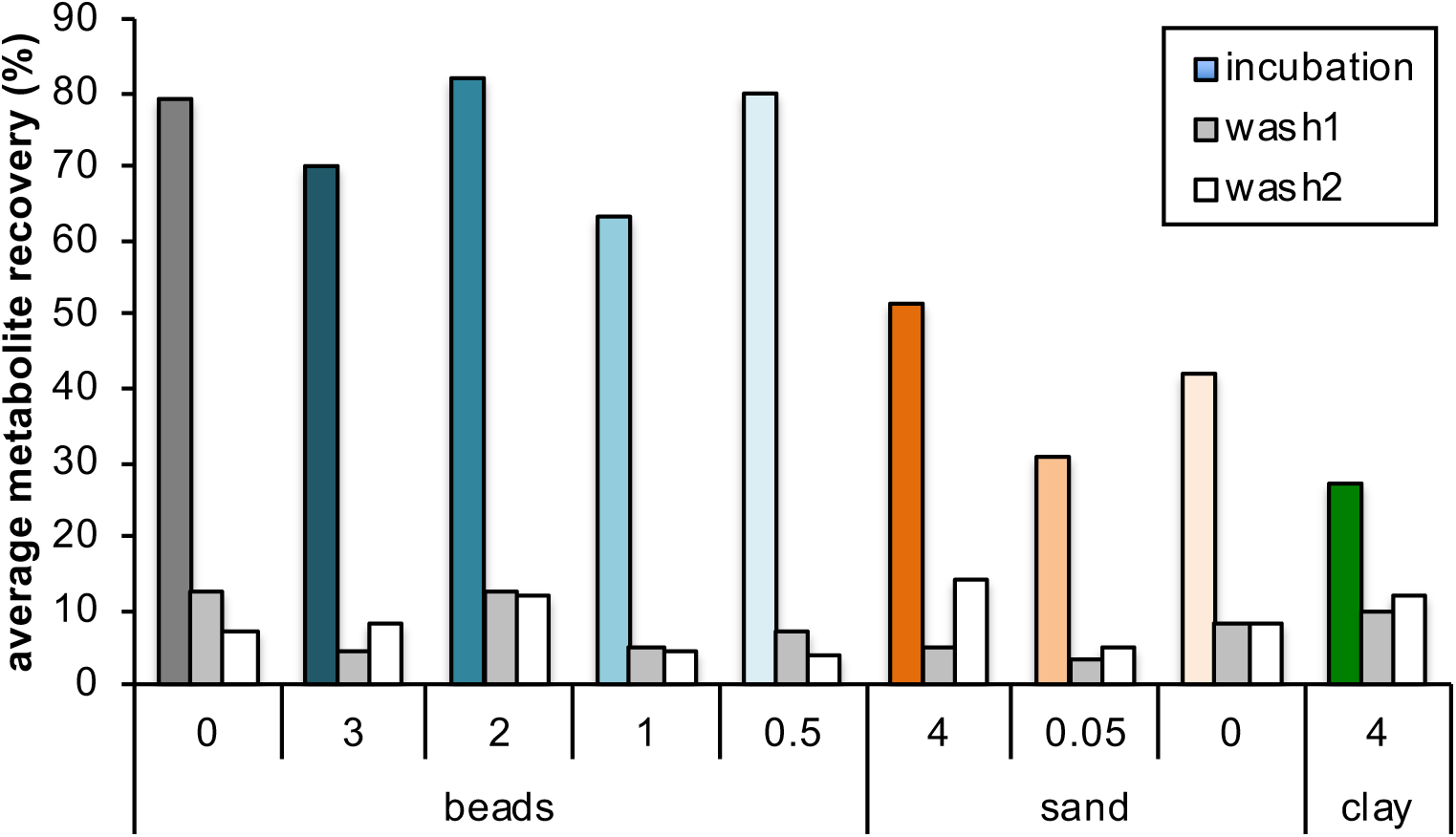
Metabolite recovery from substrates. Average defined medium metabolite recovery from various substrates after incubation (colored bars), after a first wash (grey bars), and a second wash (white bars) compared to defined medium incubated without substrate.

**Figure S11.**
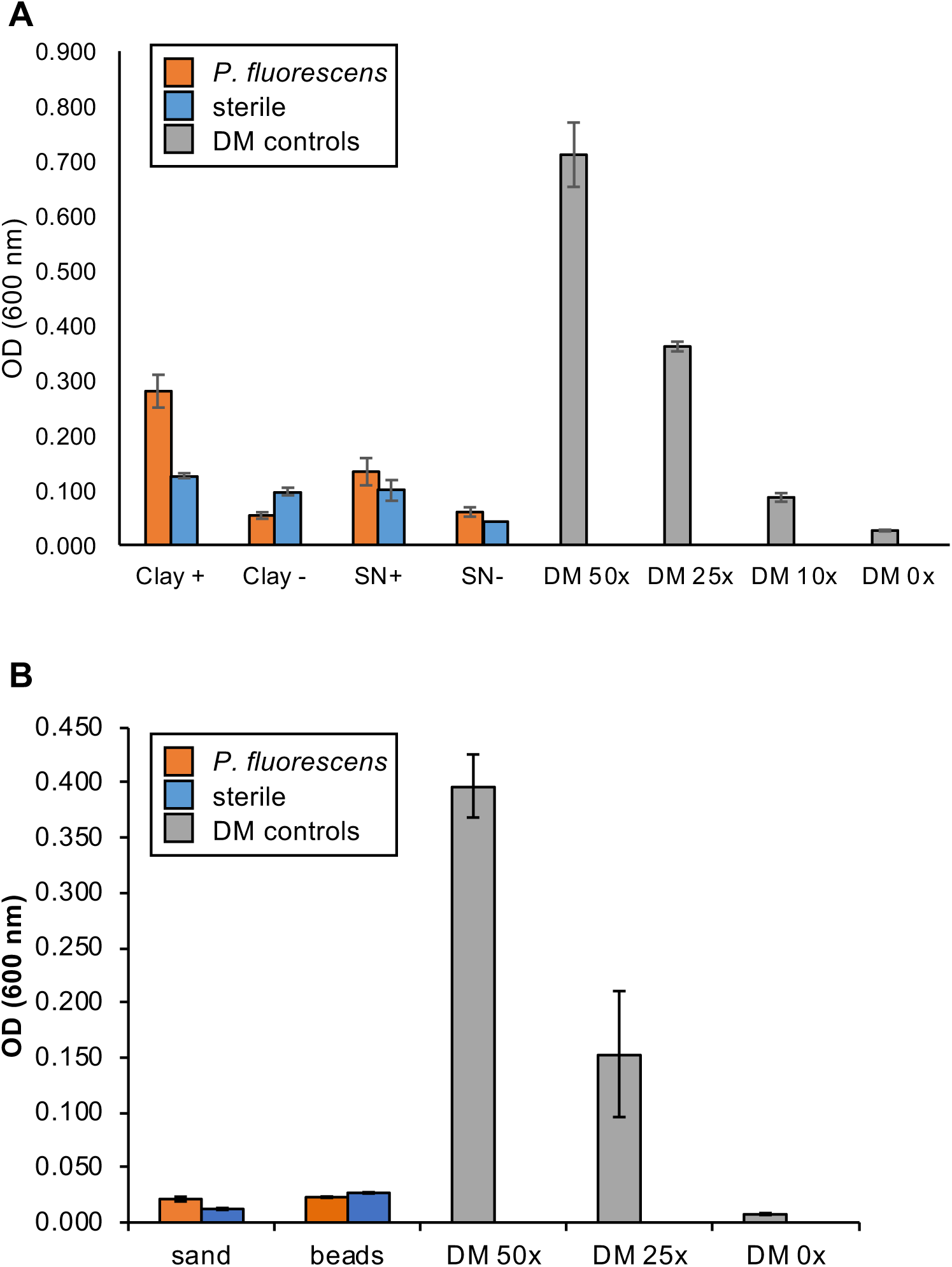
Additional data for rhizobacterium utilizes clay-adsorbed metabolites (A) Optical density (OD at 600 nm) of *P. fluorescens* (orange bars) grown in clay pre-incubated with 50x defined medium (DM, Clay +) or 0x DM (Clay -), or in supernatant of clay pre-incubated with 50x DM (SN +) or 0x DM (SN -). Sterile controls (blue bars) indicate substrate background (no bacteria control). Data normalized by substrate background is displayed in Fig. 6. **(B)** OD of *P. fluorescens* grown in 4 mm sand and 3 mm glass beads pre-incubated with 50x DM (orange) or 0x DM (blue). *P. fluorescens* growth in different concentrations of DM without substrate are given as comparison. All ODs are means ± SEM, n = 3.

**Table S1.** Diffusion Parameters for Glass Bead Substrates of Varying Sizes. Diffusion parameters calculated based on Congo Red diffusion through a column filled with glass beads of varying sizes. Values for 250 µm and 10 µm are theoretical values calculated from an exponential trend line fit.

| Bead size<br>(mm) | Diffusion rate<br>(cm h<sup>-1</sup>) | Diffusion coefficient<br>(cm<sup>2</sup> h<sup>-1</sup>) |
| --- | --- | --- |
| 3 | 21.6 | 38.9 |
| 2 | 4.48 | 8.07 |
| 1 | 1.56 | 2.50 |
| 0.5 | 1.02 | 1.08 |
| 0.25 | 0.541 | 0.718 |
| 0.005 | 0.391 | 0.504 |

**Table S2.** Metabolite data.

## Acknowledgements

The authors would like to thank Sarah Brecht and Matthew Chu for root morphology quantification, Nicholas Saichek for MALDI imaging support, and Marco Voltolini (LBNL) for X-Ray diffraction analyses. This work was part of the Microbial Community Analysis and Functional Evaluation in Soils Program at Lawrence Berkeley National Laboratory with funding from the Office of Biological & Environmental Research of the U.S. Department of Energy under contract DE-AC02-05CH11231 at the U.S. Department of Energy Joint Genome Institute, a DOE Office of Science User Facility. The research used resources of the National Energy Research Scientific Computing Center, a DOE Office of Science User Facility supported by the Office of Science of the U.S. Department of Energy under contract number DE-AC02-05CH11231. In addition, J.S. was supported by a NSF grant to University of California Berkeley (NSF Proposal 1617020).

